# Post-secretory synthesis of a natural analog of iron-gall ink in the black nectar of *Melianthus* spp.

**DOI:** 10.1101/2022.12.20.521234

**Authors:** Evin T. Magner, Rahul Roy, Katrina Freund Saxhaug, Amod Zambre, Kaitlyn Bruns, Emilie C. Snell-Rood, Marshall Hampton, Adrian D. Hegeman, Clay J. Carter

## Abstract

The black nectar of *Melianthus* flowers is thought to serve as a visual attractant to pollinators, but the chemical identity and synthesis of the black pigment are unknown. Here we report that the black nectar contains a natural analog of iron-gall ink, which humans have used since medieval times. Specifically, dark black nectar at anthesis contains high levels of ellagic acid and iron; synthetic solutions of ellagic acid and iron(III) recapitulate the black color of the nectar. Conversely, lightly colored nectars before and after anthesis contain significantly lower levels of ellagic acid and iron, but higher levels of gallic acid. We then explored the possibility of post-secretory synthesis of ellagic acid from gallic acid. Indeed, *Melianthus* nectar contains a peroxidase that oxidizes gallic acid to form ellagic acid. Reactions containing the nectar peroxidase, gallic acid, hydrogen peroxide, and iron can fully recreate the black color of the nectar. Visual modeling indicates that the black color is both visible and conspicuous to birds within the context of the flower. In summary, the black nectar of *Melianthus* is derived from an ellagic acid-Fe complex analogous to iron-gall ink and is likely involved in the attraction of passerine bird pollinators.

## INTRODUCTION

The chemical diversity of the plant kingdom is vast when compared to that of animals and microbes (Lautie *et al*., 2020). This diversity likely stems from plants’ sessile nature and the associated limitations in responding to variable environmental conditions (Knudsen *et al*., 2018). Similarly, interactions with other organisms have driven a large portion of the chemodiversity found in plants (Defossez *et al*., 2021). For instance, some plant specialized metabolites are antimicrobial (Celedon & Bohlmann, 2019), whereas others, like volatiles and colored compounds, are often involved in pollinator attraction (Bouwmeester *et al*., 2019; Fairnie *et al*., 2022).

One understudied area of plant chemical diversity is that of nectar. Plants must both attract and reward pollinators to entice repeated visitation to maximize reproductive success (Heinrich & Raven, 1972). The main reward plants offer prospective pollinators is floral nectar, which can vary widely in presentation, volume, and chemical composition across species (Roy *et al*., 2017). Nectars are always dominated by sugars (~90% of total dry wt on average) but can be chemically complex, including additional solutes like amino acids, alkaloids, flavonoids, proteins, and organic and inorganic ions (Roy *et al*., 2017). These seemingly minor components can have outsized effects on plant-pollinator and plant-microbe interactions. One example is that of the amino acid proline, which accumulates to high levels (up to ~60 mM) in the nectars of some species (Carter *et al*., 2006; Roy *et al*., 2022) and appears to play a role in pollinator preference due to a role in the metabolism of insect flight muscles (Teulier *et al*., 2016). Some nectars also contain antimicrobial metabolites and proteins to prevent microbial growth and infection (Carter, C & Thornburg, RW, 2004; Gonzalez-Teuber *et al*., 2009; Gonzalez-Teuber *et al*., 2010; Schmitt *et al*., 2018; Kurilla *et al*., 2019). In rare instances, nectars can also be colored, which is a visible cue to pollinators as a signal of the presence of an ‘honest’ reward (i.e., that a reward is truly present) (*Hansen et al*., 2007).

*Melianthus* is a genus of eight species of shrubs endemic to southern Africa (Linder *et al*., 2006) but also have widespread use as ornamentals elsewhere (Eloff *et al*., 2017). The flowers of *Melianthus minor* (syn. *elongatus*) have conspicuous red petals and produce an intensely black-colored nectar (e.g., Fig. 1); other species in the genus produce either black or tan nectar, in contrast to the non-colored nectar of the most closely related genus, *Bersama* (Decraene *et al*., 2001; Henning, 2003; Linder *et al*., 2006). The nectar’s black coloration is suggested to serve as a visual cue to passerine bird pollinators (Henning, 2003; Hansen *et al*., 2007), but the chemical nature and true function of the black pigment are unknown. Our research aimed to identify the black pigment, routes of its synthesis, and explore the nectar’s biological function as a conspicuous cue to avian pollinators.

**Fig. 1.**
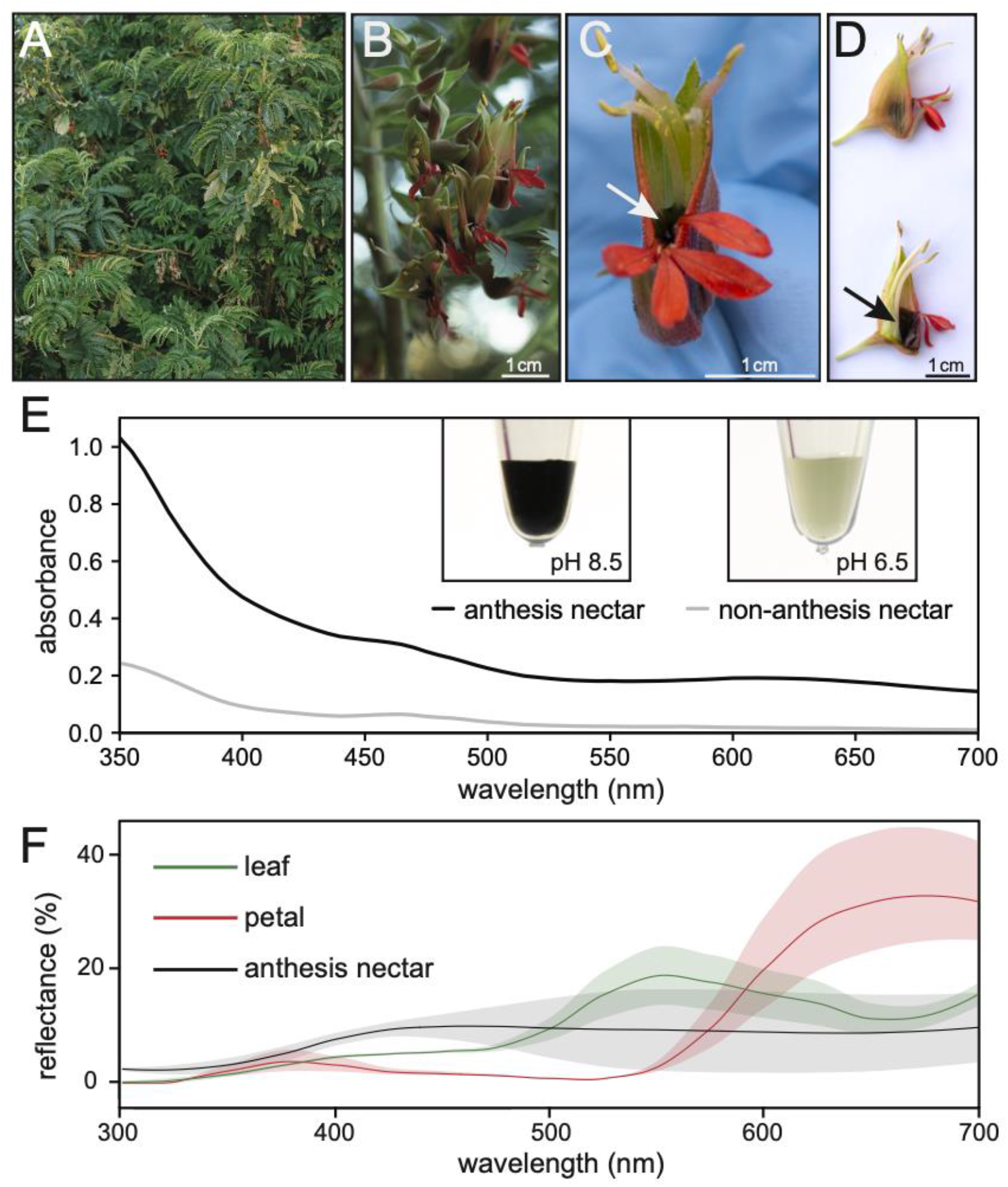
*Melianthus minor* floral morphology and nectar characteristics. (A) *M. minor* growth form that conceals the majority of inflorescences within the bush. (B) *M. minor* inflorescence, with flowers at varying stages, including pre-anthesis, anthesis, and post-anthesis. (C) An individual flower at anthesis showing the black nectar (arrow) within the fused sepals and surrounded by red petals. (D) Lateral view of a pre-anthesis flower (top, anthers not dehisced) and anthesis flower (bottom, partial longitudinal section), showing the position of the nectar (arrow) inside of the flower. (E) Absorbance spectra of anthesis (dark hue) and non-anthesis (light hue) nectar diluted 1:20 in diH_2_O. Inset images are of undiluted samples with the pH noted. (F) Reflectance spectra of the flower, leaf, and anthesis nectar.

## MATERIALS AND METHODS

### Plant materials & collection

*Melianthus minor* (also known as *M. elongatus*) and *M. comosus* plants were grown at the University of California Davis Arboretum and Public Garden at the University of California Davis campus in Davis, California, USA. Plants were grown and maintained outdoors on campus and received yearly maintenance of pruning and the application of fertilizer as needed.

From 26 January 26 2022 to 1 February 2022, leaf, flower, nectary, and nectar samples were collected from the University of California Davis Arboretum and Public Garden. Collections took place each day between 10:00 and 17:00. Plant tissue were manually collected and were either placed immediately in RNAlater stabilizing solution (Thermo Fisher AM7020) or on dry ice before being stored at −80°C. Additional nectar samples from *M. pectinatus* and *M. villosus* were collected in the field in South Africa.

### Description of nectar and nectary stages

Nectar samples were collected and sorted into two groups: anthesis and non-anthesis nectar. Nectar produced following anther dehiscence and up to two days after was categorized as anthesis nectar. Peak nectar production occurred for two days following the dehiscence of anthers. Nectar produced prior to anther dehiscence was classified as non-anthesis. Nectar collected from flowers 3+ days following anthesis was also categorized as non-anthesis when transparent (i.e., lightly colored relative to anthesis nectar) to any degree. These light-hued, post-anthesis nectar samples were additionally pooled with pre-anthesis nectar samples. Together, the pre-anthesis and post-anthesis light-hued nectars constituted the ‘non-anthesis’ time point. However, the pooled samples were dominated by the pre-anthesis time point.

Nectary samples were divided into stages determined by the state of the anther, nectar production, and the development of each flower on the inflorescence (Fig. S1). Nectaries at the pre-secretory stage are described as the anthers being neither exposed nor dehisced. The petals remain fused and are not present past the sepals, which are still fused at this point. The secretory stage nectaries were collected from flowers whose anthers have dehisced, the petals have emerged from the sepals, and the flower is visibly open. Furthermore, nectar is actively being produced and dark black in color. The post-secretory stage nectaries were from flowers that had ceased to produce nectar, with anthers being dehisced for more than 3 days, and in some cases, the fruit had begun to develop. The primary distinction between stages is the presence or absence of nectar for the RNAseq analyses described next.

### Transcriptomic analyses

RNA was isolated from pre-secretory, secretory, and post-secretory nectaries (stages described above), as well as leaves, with the RNAqueous Micro Kit (Invitrogen cat. no. AM1931). The total RNA was treated with the TURBO DNA-free kit (Invitrogen Cat# AM1907) according to the manufacturer’s protocol and submitted to the University of Minnesota Genomics Center for mRNA isolation, barcoded library creation and sequencing via NovaSeq 6000 using via paired-end 150 bp runs using rapid chemistry.

Trinity version 2.12.0 (Grabherr *et al*., 2011) was used to assemble 462,684 contigs from the reads de novo. Predicted protein sequences for all contigs were computed using Trinity’s Transdecoder (version 5.5). Normalized read counts for the assembled contigs were computed using kallisto (version 0.46.1) (Bray *et al*., 2016), and then upper-quartile normalized to correct for variable rRNA content. The contigs were then mapped to the Araport11 *Arabidopsis thaliana* reference nucleotide and protein sequences using NCBI’s BLAST+ suite (version 2.6.0). Pairwise differential expression between sample types was tested using DESeq2 (version 1.26.0) within R (Love *et al*., 2014), using a false discovery rate cutoff of 0.05.

### pH Measurement

The pH of freshly collected anthesis and non-anthesis nectar was determined using McolorpHast 5.0 to 10.0 pH strips (EM-Reagents catalog no. 1095330001). Later, a Thermo Scientific Orion Star A214 pH Benchtop Meter was used to confirm the pH of the pooled anthesis nectar.

### UV/Visible absorbance spectroscopy

The UV/visible light absorbance of nectar samples, synthetic pigments, and substrates was measured employing a BioTek Powerwave HT 96-well plate reader, an Implen NanoPhotometer Pearl, or a Beckman DU 520 General Purpose UV/Vis Spectrophotometer.

### Reflectance spectroscopy

We used a UV-VIS spectrophotometer [OceanOptic JAZ-A2474 with a pulsed xenon (PX) lamp] to collect reflectance spectra of *M. minor* petals and nectar (N = 3 flowers). The triggering rate of the PX lamp was 10ms, and the boxcar was set to 5. The spectral probe was held at an angle of 45° against *M. minor* flowers. The flowers were placed on black velvet paper to minimize measurement errors. For nectar, 100 μL nectar was pipetted on a fresh Kimwipes tissue paper and allowed to dry. Reflectance data of dried nectar was then recorded using the same protocol for measuring the reflectance of flowers. All measurements were corrected against white and black reflectance standards. Collected spectral data were imported in R, smoothened (α = 0.35), and negative values were fixed to ‘0’ using the ‘fixneg’ function using ‘pavo2.0’ R package. These corrected and smoothened spectra (Fig. 1D) were used for subsequent visual modeling (Fig.7).

**Fig. 2.**
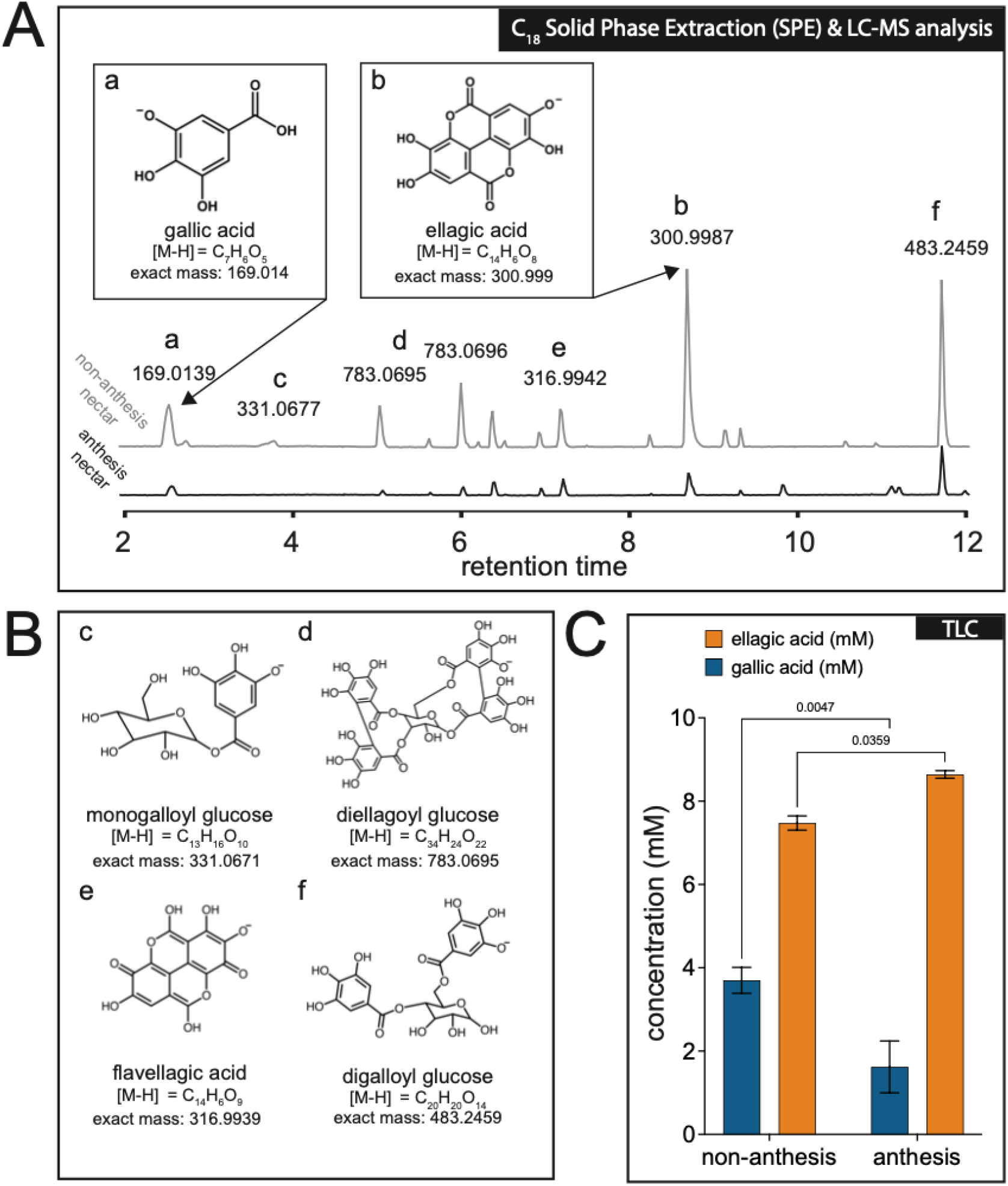
*M. minor* nectar contains high concentrations of gallic and ellagic acid. (A) LC-MS analysis revealed gallic acid and ellagic acid to be major components in nectar samples. (B) Structures, formulas, and masses corresponding to other significant peaks structurally related to gallic and ellagic acid. The letters correspond to the peaks in (A). The true positions of gallic acid on the glucose of mono- and digalloyl glucose are unknown. (C) Concentrations of gallic and ellagic acid in non-anthesis and anthesis nectars as determined by TLC (p-values derived from two-way ANOVA; N = 3).

**Fig. 3.**
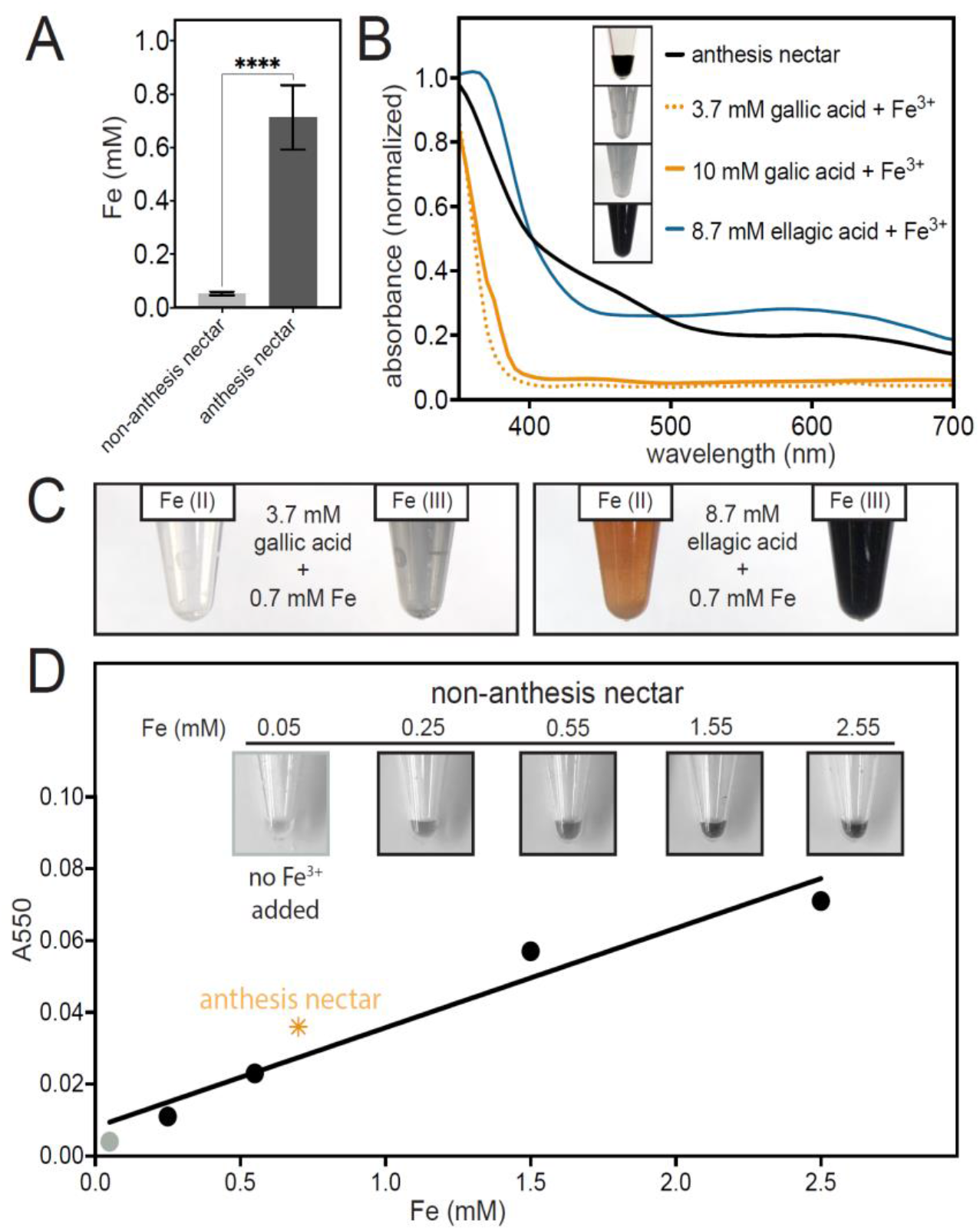
*M. minor* nectar contains high concentrations of iron, which is required for black pigment formation. (A) Iron content of *M. minor* nectar from anthesis and non-anthesis flowers (two-tailed t-test p<0.0001). (B) Absorbance spectra of biologically relevant concentrations of ellagic acid (8.7 mM) and gallic acid (at 3.7 mM and 10 mM) after addition of 0.7 mM FeCl_3_. (C) Visual appearance of tubes after addition of 0.7 mM FeCl_3_ [Fe(III)] or FeSO4 [Fe(II)] to 3.7 mM gallic acid or 8.7 mM ellagic acid. (D) Addition of varying concentrations of FeCl_3_ to non-anthesis nectar (grey box and grey point have no Fe added). Insets show successive images of the same non-anthesis nectar sample with increasing final concentrations of iron. Absorbance as a function of [Fe] was read at 550 nm, a region of the spectrum with minimal individual absorbance of Fe(III), gallic acid, and ellagic acid. The R^2^ of the line is 0.9382. The absorbance of a typical anthesis nectar (diluted 1:20) at 550 nm plotted versus its associated [Fe] of 0.7 mM is shown with the orange asterisk.

**Fig. 4.**
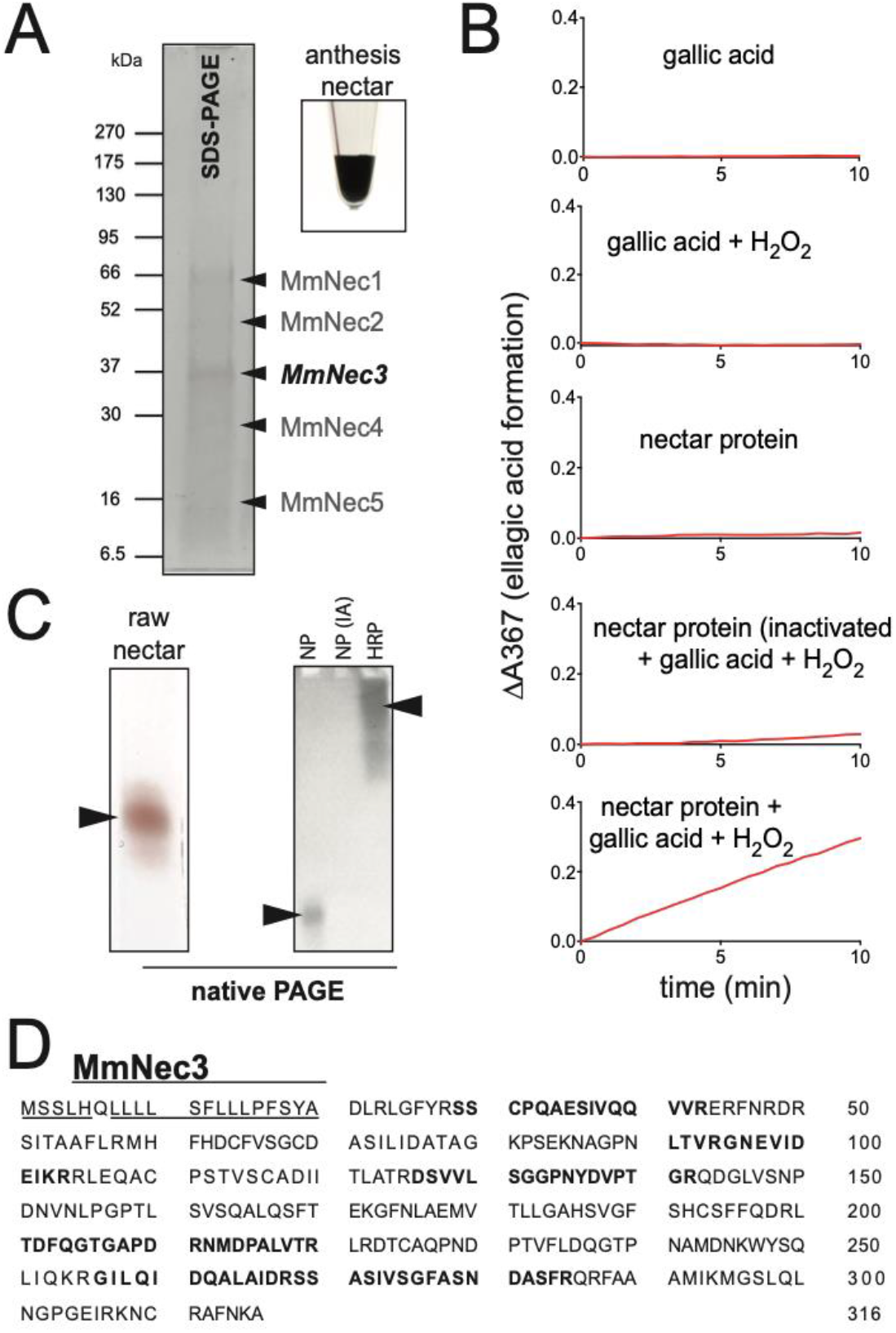
A nectar peroxidase can oxidize gallic acid into ellagic acid. (A) SDS-PAGE analysis (4-20%) of 16 μL of raw *M. minor* nectar stained with Coomassie Blue. Arrowhead indicates the location of MmNec3, which was identified as a putative peroxidase by proteomic analysis. (B) Peroxidase activity of concentrated total nectar protein in 1x TBS pH 9.0. Individual assays contained the components shown in each subpanel. The change in absorbance at 367 nm (red lines) was used to detect the production of ellagic acid. Total nectar protein was inactivated with Proteinase K as a negative control. (C) Native PAGE and in-gel peroxidase activity of nectar proteins (NP) using guaiacol (left) or gallic acid (right) as substrates. Inactivated nectar protein [NP-IA] and horseradish peroxidase (HRP) was used as negative and positive controls for gallic acid oxidation, respectively. FeCl_3_ was used for color development of ellagic acid. (D) Translated sequence of MmNec3 (NCBI accession # pending) with peptides identified from the activity band with guaiacol from panel C bolded. The underlined region corresponds to a predicted N-terminal signal peptide.

**Fig. 5.**
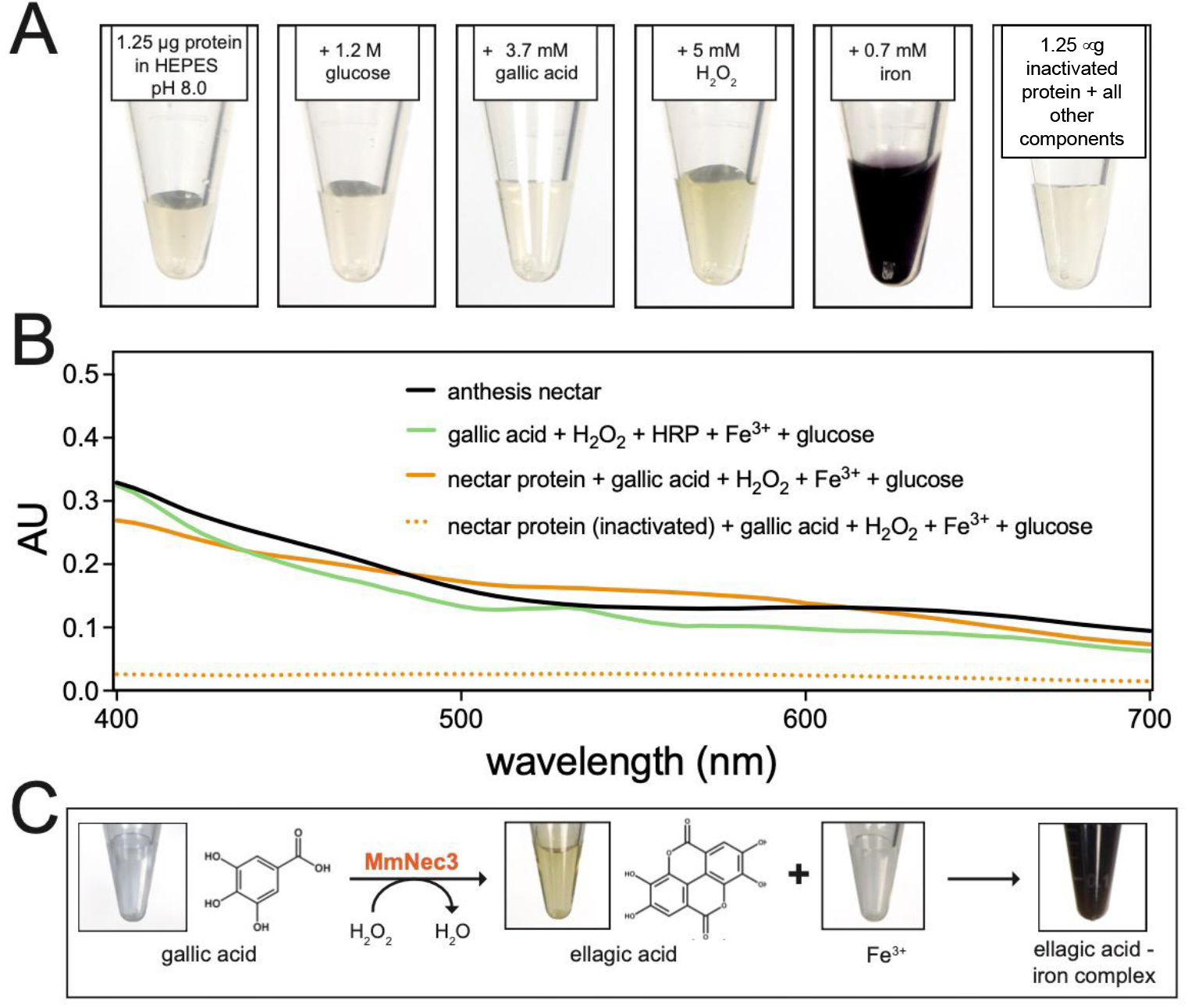
In vitro recapitulation of black pigment formation. (A) Total nectar protein (1.25 μg) in 25 mM HEPES pH 8.0 with nectar components sequentially added to the final concentrations shown in each inset. Images show the same tube after components were added sequentially from left to right. A negative control reaction containing inactivated nectar protein (Proteinase K treated) and all other components is shown on the far right. (B) Normalized absorbance spectra of anthesis and synthetic nectars. The spectrum from the reaction using total nectar protein from panel A is shown, as well as reactions using HRP as a positive control and Proteinase K-inactivated total nectar protein as a negative control. (C) Proposed reaction sequence between gallic acid, H_2_O_2_, and MmNec3 to yield ellagic acid, which then complexes with Fe(III) to yield a black color.

**Fig. 6.**
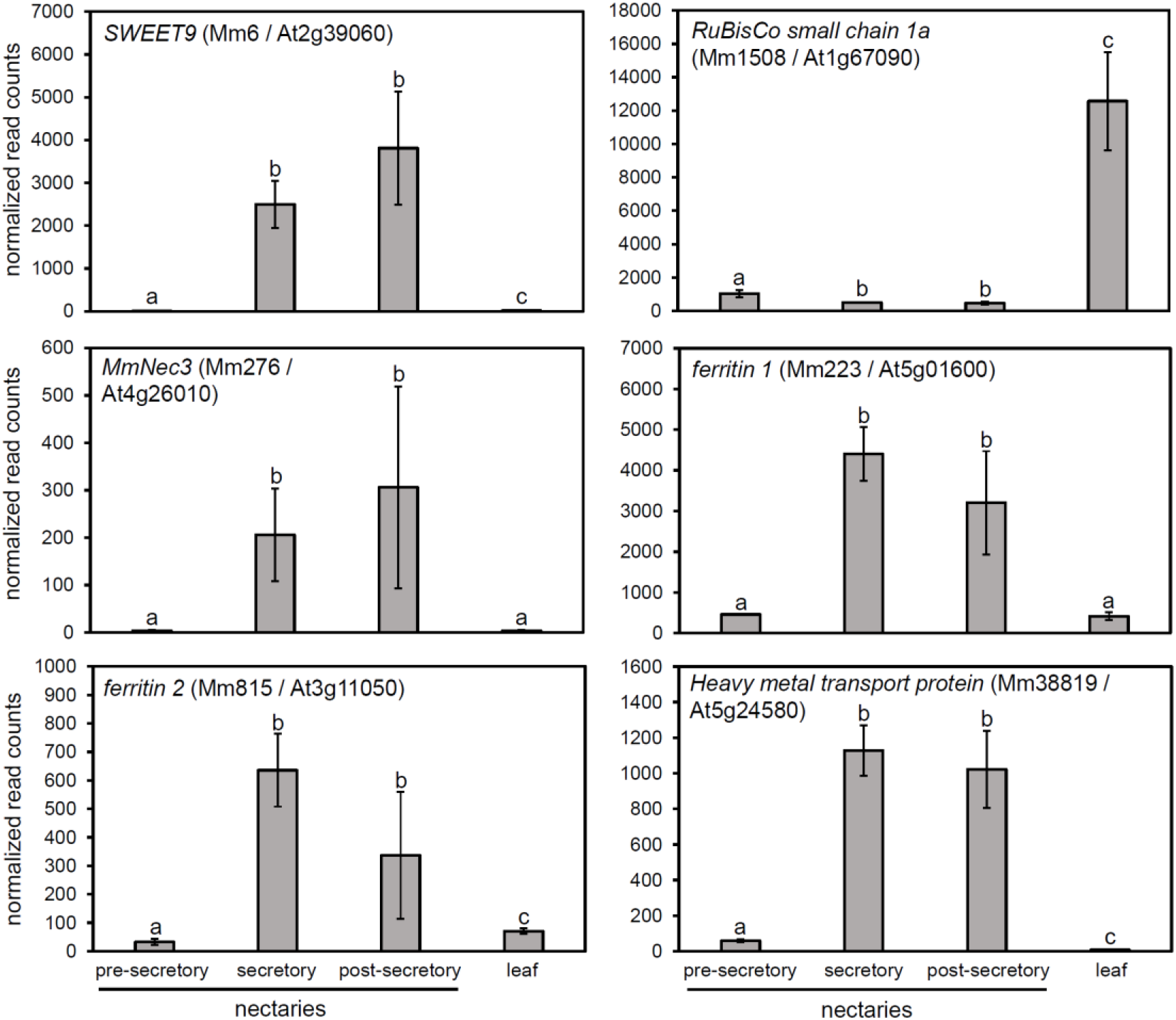
Expression of MmNec3, the nectar peroxidase, is strongly induced during nectar secretion. Normalized read counts derived from RNA-seq analysis are shown for select genes in pre-secretory, secretory, and post-secretory nectaries, as well as leaves. Each panel indicates the putative identity along with the contig number (e.g., Mm6) and top Arabidopsis hit (e.g., At2g39060). Triplicate samples were used for all tissues except for pre-secretory nectaries, which only had two samples. Error bars represent standard deviation and different letters represent significant differences in gene expression as determined by DEseq analysis. Summarized data are shown in Supplemental File 1 and raw data is available on NCBI SRA via accession # (pending).

**Fig. 7.**
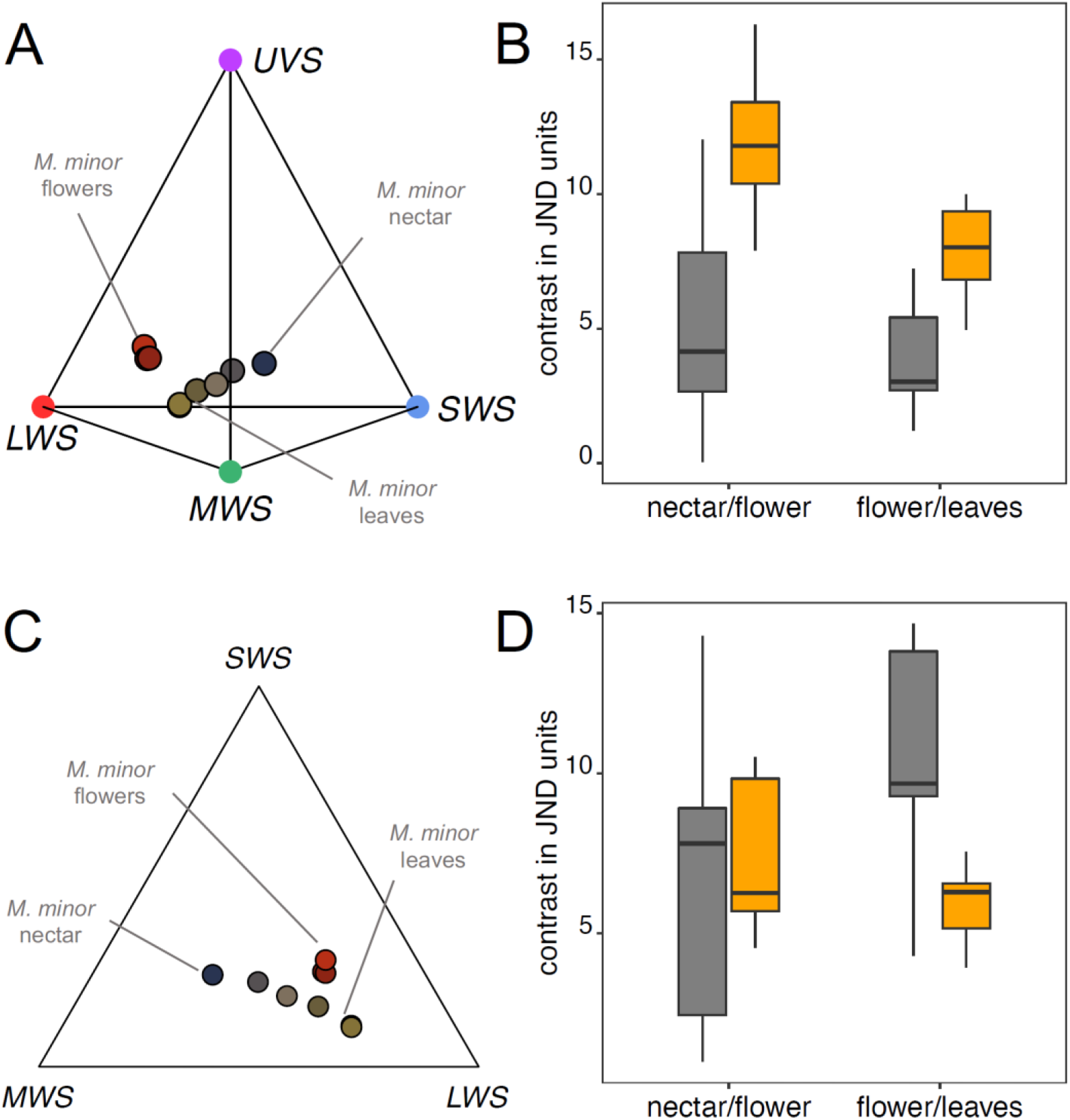
The black nectar is the most conspicuous signal for birds but not bees. (A) Tetraplot showing *M. minor* reflectance data projected onto *Sturnus vulgaris* visual space. The vertices of the tetrahedron correspond to four different photoreceptors (B) Chromatic (orange box) and achromatic contrasts (grey box) of petals and the natural plant interior background and nectar and the adjacent petals for *M. minor* using *S. vulgaris* photoreceptor sensitivities. (C) *Melianthus minor* reflectance data projected onto *Apis* bee visual space. (D) Chromatic (orange box) and achromatic contrasts (grey box) of petals and the natural plant interior background and nectar and the adjacent petals for *M. minor* using *Apis* photoreceptor sensitivities. Contrasts are expressed in units of JND (just noticeable differences), and the higher the value, the more conspicuous the color should appear to pollinators.

### Nectar proteomics

For the initial identification of MmNec3, 16 μL of *M. minor* nectar was electrophoresed on a 4-20% Mini-PROTEAN TGX gel under denaturing conditions, and PAGE-Blue (Thermo Fisher Scientific catalog no. 24620) was used to stain the gel. Excised protein bands were submitted for identification to the University of Minnesota’s Center for Mass Spectrometry and Proteomics Facility as previously described by our group (Roy *et al*., 2022).

### Purification of total nectar protein

Raw nectar samples were first centrifuged at 13,300 x g for 10 minutes at 4°C to remove any large debris. Then 400 μL of the clarified nectar was subsequently centrifuged through a 10,000 molecular weight cut-off (MWCO) Amicon Ultra-0.5 Centrifugal Filter (Millipore Sigma SKU UFC5010) for 10 min at 13,300 × g at 4°C. The retentate was diluted back up to 400 μL in 50 mM HEPES (pH 8.0) and centrifuged two more times, increasing the centrifuge time to 20-30 minutes. The samples were then washed a total of three times in 25 mM HEPES (pH 8.0). Next, the nectar samples were washed an additional three times using 10 mM HEPES (pH 8.0). The final wash prior to recovery used 25 mM HEPES (pH 8.0). The Bradford assay determined the total protein concentration in nectar samples. Concentrated nectar proteins were evaluated using denaturing 4-20% SDS-PAGE gels stained with PAGE-Blue (Thermo Fisher #24620).

### Nectar sugar determination

Total sugar content of nectar samples was determined using a refractometer (Bolten *et al*., 1979). Accurate estimations were obtained by comparing the relative (%) sugar content to that of sterile water. The percent sugar content was converted to molarity, and thus molarity is noted in the following work. A colorimetric invertase assay, as previously described, was used to determine the composition of soluble sugars (Minami *et al*., 2021).

### Liquid Chromatography-High Resolution Mass Spectrometry (LC-HRMS) analysis of nectar metabolites

Nectar samples were first thawed on ice and then centrifuged at 13,300 x g for 5 minutes at 4°C. The supernatant was transferred to a new tube and pigments were isolated from the raw nectar using an adapted protocol using ZipTipC18 pipette tips as previously described (Roy *et al*., 2022). The eluate was then transferred into LC-MS autosampler vials and 1 μL was injected (via autosampler) onto a reversed-phase C18 HSS T3 1.8 μm particle size, 2.1 x 100 mm column (Waters, Milford, MA, USA). Column temperature was 40°C and solvent flow rate 0.40 mL/min. A 29-minute linear gradient using mobile phases A: 0.1% formic acid in water and B: 0.1% formic acid in acetonitrile was run according to the following gradient 29 minute elution profile: initial, 2% B; 2 min, 2% B; 25 min, 98% B; 1 min, 98% B, 1 min 2% B. UV-vis data was collected from 200-600nm for duration of gradient. The following MS conditions were used: full scan mass scan range: 125-1800 *m/z*, resolution: 70,000, data type: profile, desolvation temperature 350°C, capillary voltage: 3800 V (+), 4000 V (-). Xcalibur™ software version 2.1 (Thermo Scientific, Waltham, MA, USA) was used to record and visualize the chromatograms and spectra.

### Gallic acid and ellagic acid quantification via TLC

Gallic acid and ellagic acid concentrations were determined using a previously published procedure using thin-layer chromatography (TLC) (Kamel *et al*., 1977; Wagh, 2010). Standards of gallic acid and ellagic acid of varying concentrations (0 - 10 mM) were used to calculate both concentrations in triplicates of non-anthesis and anthesis nectar samples. The stationary phase was silica on glass, pre-treated in a 100°C vacuum oven overnight. The mobile phase consisted of a solution of 5 parts chloroform, 4 parts ethyl acetate, and 1 part formic acid. 1 μL of standards and samples were spotted on the TLC baseline and placed in a chamber containing 7 mL of the mobile phase. Images were first turned into a grey-scale and analyzed by ImageJ. Areas of the same size and shape were used to construct a standard curve, from which concentrations samples of non-anthesis and anthesis nectar could be calculated from the mean grey value.

### Iron quantification

Total iron was measured using a previously published ferrozine-based colorimetric assay (Riemer *et al*., 2004). Nectar samples [anthesis n=15, non-anthesis n=14] were first diluted 1:10 using sterile water. Iron standards of varying concentrations [0 mM – 0.1 mM Fe] were created using FeCl_3_. Iron content was averaged for anthesis and non-anthesis, and a two-tailed Student’s t-test was used to evaluate the significance of any difference.

### In-gel enzymatic assays (zymography)

Two types of in-gel activity assays were used to detect peroxidase activity in the nectar. The first method used guaiacol as a chromogenic substrate as previously described (Wilkesman *et al*., 2014) with a single replicate consisting of 16 μL of raw nectar electrophoresed on a non-denaturing 4-20% native polyacrylamide gel. A second method for the in-gel detection of peroxidase activity relied on the production of ellagic acid from gallic acid. Nectar samples were treated the same as for the guaiacol-based assay described above, except after electrophoresis and rinsing in diH2O, the gel was placed in a substrate solution containing 20 mM gallic acid in 10 mM sodium phosphate buffer pH 7.4 for 30 minutes. The gel was then briefly rinsed with diH2O and placed back into 10 mM sodium phosphate buffer pH 7.4. To this, we added 100 μL of a developing solution containing equal parts 3% H_2_O_2_ and 100 μM FeCl_3_. A black activity band developed immediately in areas containing peroxidase activity due to the conversion of gallic acid into ellagic acid. Positive control lanes contained 1 U of HRP.

#### Spectrophotometric determination of conversion of gallic acid to ellagic acid

Reactions contained total nectar protein (1.25 μg), 20 mM gallic acid, and 1 mM H_2_O_2_. The absorbance at 367 nm (the major absorbance peak for ellagic acid) was then measured over 10 minutes. Blanks omitted the addition of nectar protein. For negative controls, either nectar proteins were first inactivated by Proteinase K treatment (Thermo Fisher Scientific catalog no. 25530049) or H_2_O_2_ was omitted.

### Visual modeling

We used published cone sensitivities for European starling *Sturnus vulgaris* (UVS ƛ_max_ = 362, SWS ƛ_max_= 449, MWS ƛ_max_= 504, LWS ƛ_max_= 563; ratio of UVS: SWS: MWS: LWS = 1:2:2:4; from (Hart *et al*., 1998), which is a known pollinator of *M. minor*. For the bee visual model, we used photoreceptor sensitivities of the honey bee *Apis mellifera* (SWS ƛ_max_ = 344, MWS ƛ_max_= 436, LWS ƛ_max_= 544; sensmodel=‘apis’ in pavo2.0). For both *S. vulgaris* and *A. mellifera*, we used RNL models (Receptor-Noise Limited models) and set the irradiance to bright daylight (‘D65’). We also applied von Kries transformation to account for light adaptation (Vorobyev *et al*., 2001). *M. minor* reflectance spectra (see above) were projected onto the *S. vulgaris* and *A. mellifera* visual spaces. *Sturnus vulgaris* has four distinct cone receptors whereas *A. mellifera* has only three distinct photoreceptors. Therefore, the visual space of *S. vulgaris* is represented as a tetrahedron while that of *A. mellifera* is represented as a triangle with each vertex corresponding to a specific cone. Points within the visual space represent the reflectance of flowers and nectar, and their location within visual space estimated by stimulation of each cone type.

To examine how conspicuous nectar and flowers appear to birds, we used our visual models to calculate chromatic contrast (differences in hue and chroma) and achromatic contrast (difference in luminance) of flowers and nectar against their likely natural backgrounds as viewed by the pollinators. Contrasts values are a function of the object of interest (*M. minor* nectar and flowers) and the background against which they are viewed. Higher contrast values indicate greater conspicuousness and are generally associated with greater pollinator attraction. To calculate the contrast of *M. minor* nectar, we used the flower as a background, whereas for flowers, we used *M. minor* leaves as the background.

## RESULTS

### Nectar presentation and characteristics

Our study focused on *Melianthus minor* (also known as *M. elongatus*) grown on the campus of the University of California Davis as part of its arboretum collection and as an ornamental. *M. minor* inflorescences tend to be located on the interior of the shrub (Fig. 1A), with individual flowers of a single inflorescence maturing at different times (Fig. 1B). Open flowers have red-colored petals and produce copious nectar (Fig. 1C). Nectar secretion begins after the petals open but before anthesis (i.e., pre-anthesis, as defined by the presence of non-dehisced anthers) with maximal secretion occurring at anthesis and continuing for approximately two days. By three days after anthesis, nectar volume begins to significantly decline (i.e., post-anthesis).

Both pre- and post-anthesis nectars (collectively referred to as non-anthesis nectar hereafter) are lightly colored and have an acidic pH, whereas the nectar at anthesis is intensely black and has an alkaline pH (Fig. 1E). The black nectar has nearly uniform absorbance from 400-700 nm, with much higher absorbance in the UV region under 400 nm. Conversely, non-anthesis nectar has relatively little absorbance over this same spectrum. Not surprisingly, the black anthesis nectar has very little reflectance from 300-700 nm, with leaves and petals having highest reflectance in the green and red regions of the visible spectrum, respectively (Fig. 1F).

### *Melianthus* nectar is rich in gallic and ellagic acid

To evaluate what nectar compounds might be responsible for the black coloration, nonpolar nectar solutes were partially purified by solid phase extraction (i.e., ZipTip with C18 resin) and subjected to liquid chromatography mass spectrometry (LC-MS). Polyphenols are particularly prevalent in both non-anthesis and anthesis nectars, including free (Fig. 2A) and glucosylated (Fig. 2B) forms of gallic acid (i.e., gallotannins). Up to five gallic acid moieties can be directly conjugated to glucose in plants (i.e., mono-, di-, tri-, tetra-, and pentagalloyl glucose), with head-to-tail additions of gallic acid being possible to form decagalloyl glucose, also known as tannic acid (Grundhofer *et al*., 2001). These variable galloyl glucose molecules are collectively known as gallotannins (Grundhofer *et al*., 2001). Mono-, di-, and tri-galloyl glucose are present in *M. minor* nectar (Table S1), suggesting that gallic acid may be actively cleaved off of glucose in a post-secretory manner. Ellagic acid, formed from the oxidative dimerization of gallic acid, appears to be the most abundant polyphenol in *Melianthus* nectars (Fig. 2A, Fig. 2C, and Fig. S3). We also observed dimers of apparently adjacent galloyl residues attached to glucose (i.e., ellagitanins; compound b in Fig. 2B) which, upon hydrolysis from glucose and lactonization, can form ellagic acid (Grundhofer *et al*., 2001). Flavellagic acid, a derivative of ellagic acid, was also found in both anthesis and non-anthesis nectars. Similar patterns of gallic acid, ellagic acid, and related tannins and polyphenols were observed in the nectars of other *Melianthus* species, including *M. comosus, M. pectinatus, and M. villosus* (Fig. S3, Table S1).

The high levels of gallic acid in the nectar suggested its coloration might be derived from a natural analog of iron-gall ink. This ink was used extensively from medieval times up through the middle of the 20^th^ century, with recipes often involving the addition of vitriol [ferrous sulfate (FeSO4)] to extracts of ground oak galls, which are high in gallic acid and hydrolysable tannins (Melo *et al*., 2022). More modern recipes for iron-gall ink even explicitly call for gallic acid and tannic acid (decagalloyl glucose) as ingredients (Wailes, 1935). To test the hypothesis that *Melianthus* nectar is a natural analog of iron-gall ink, we evaluated the gallic and ellagic acid content of anthesis and non-anthesis nectars by thin-layer chromatography (Fig. 2C and Fig. S2). The dark anthesis nectar has significantly less gallic acid than the lighter colored non-anthesis nectar (~1.5 mM in dark vs. ~3.7 mM in light), with the reverse being true for ellagic acid (~8.6 mM in anthesis vs. ~7.3 mM in non-anthesis nectar). While there are statistically significant differences in total gallic and ellagic acid in the anthesis vs. non-anthesis nectars, we believed these differences unlikely to be solely responsible for the stark discrepancy in subjective color intensity.

### Iron and ellagic acid are required for black pigment formation

Since the relatively small difference in gallic and ellagic acid content is unlikely responsible for the change in coloration between non-anthesis and anthesis nectars, we next examined if Fe content might play a role. Indeed, anthesis nectars contain significantly more Fe than non-anthesis nectars (Fig. 3A; ~0.7 mM Fe in anthesis vs. ~50 μM in non-anthesis). Interestingly, mixtures of biologically relevant concentrations of Fe (0.7 mM) and gallic acid (3.7 mM) at pH 8.5, the pH of anthesis nectar, produce a light grey color but do not recapitulate the black coloration of anthesis nectar, which was true at even 10 mM gallic acid (Fig. 3B). However, mixtures of ellagic acid (8.6 mM), the major polyphenol in the nectar, and FeCl_3_ (0.7 mM) readily recapitulated the black coloration of anthesis nectar (Fig. 3B). Absorbance spectra of additional ratios and mixtures of gallic acid, ellagic acid, Fe^2+^ and Fe^3+^ are shown in Fig. S4. As mentioned above, the lightly colored non-anthesis nectar is slightly acidic (pH ~6.5) and the dark anthesis nectar is alkaline (pH ~8.5). However, the pH of the nectar does not appear to influence pigment intensity, as making non-anthesis nectar more alkaline, or anthesis nectar more acidic, does not greatly impact the nectar color (Fig. S5). These results strongly suggest that (1) Fe is the limiting factor in color formation in non-anthesis nectar, (2) ellagic acid instead of gallic acid is the main color-forming agent in *Melianthus* nectar, and (3) nectar pH does not greatly impact pigment formation.

We were next interested in determining if the production of a black pigment requires ferric (Fe^3+^) or ferrous (Fe^2+^) ions in combination with either gallic or ellagic acid. Again, the addition of Fe^2+^ or Fe^3+^ to gallic acid gives either no color or a gray product (Fig. 3C). Conversely, the addition of Fe^2+^ to ellagic acid gives a reddish color, whereas Fe^3+^ and ellagic acid gives a black product of similar appearance and absorbance spectrum to anthesis nectar (Fig. 2C). To further support the idea that Fe^3+^ is the limiting factor in color formation of non-anthesis nectar, successive additions of increasing amounts of FeCl_3_ (Fe^3+^) to the same non-anthesis nectar sample leads to a concomitant and linear increase in color intensity (Fig. 3D), likely due to the high levels of pre-existing ellagic acid (Fig. 2C). These results cumulatively indicate that complexes of Fe^3+^ and ellagic acid are primarily responsible for the characteristic black color of anthesis nectar.

The observed differences of gallic and ellagic acid in non-anthesis and anthesis nectars (Fig. 2C) suggest that a dynamic process controls their accumulation. Intriguingly, the total phenolic content (ellagic + gallic acid) is roughly the same between the light and dark colored nectars, but anthesis nectar has significantly lower gallic acid and higher ellagic acid. This finding is particularly important since gallic acid is a precursor of ellagic acid (Kamel *et al*., 1977).

### A nectar peroxidase can oxidize gallic acid to form ellagic acid

We previously reported that nectar enzymes are important for the formation of a red colored alkaloid (nesocodin) in the nectar of the gecko-pollinated flowers of *Nesocodon mauritianus* (Roy *et al*., 2022). To test if the same is true for *Melianthus*, we examined its nectar proteome via SDS PAGE analysis. *M. minor* nectar contains five major proteins (MmNec1 through MmNec5), with the predominant one being MmNec3 (Fig. 4A). Since limited genomic resources exist for *Melianthus* spp. we performed RNA-seq analysis of its nectaries and leaf tissue (described in greater detail below) to generate a translated database (Files S2-S4). This predicted translatome allowed us to identify MmNec3 as a putative peroxidase via routine tryptic digest and LC-MS/MS analysis, with peptides covering 64% of the predicted mature protein being identified (Fig. S6). A putative full-length cDNA contig for MmNec3 encodes a 316 amino acid long protein with a predicted canonical 20 amino acid N-terminal signal peptide required secretion from the cell [predicted by SignalP 6.0 (Teufel *et al*., 2022)]. The mature protein (minus the signal peptide) has a theoretical isoelectric point of 7.05, a potential N-glycosylation site at Asn90, and a mature molecular weight of 32.2 kDa.

Conserved domain analysis showed that MmNec3 is a Type III peroxidase. Members of this peroxidase family, including horseradish peroxidase (HRP), contain a heme cofactor and are often associated with the cellular secretory pathway (i.e., endomembrane system and apoplast) (Shigeto & Tsutsumi, 2016). Collectively, they catalyze the oxidation of a wide range of substrates and use hydrogen peroxide (H_2_O_2_) as an electron acceptor. For instance, in 1975, a protein extracted from turnip with peroxidase activity was reported to catalyze the oxidation of gallic acid, which in turn would dimerize to form ellagic acid via an unknown reaction mechanism (Kamel *et al*., 1977). We first tested for peroxidase activity in total nectar protein (purified from other nectar components) via a spectrophotometric assay using gallic acid as a substrate and looking for ellagic acid as a product. Reactions containing gallic acid, H_2_O_2_, and total nectar protein readily generate ellagic acid (Fig. 4B), which was confirmed by thin layer chromatography (TLC; Fig. S7). Negative control reactions either lacking nectar proteins or containing inactivated nectar proteins (treated with Proteinase K) produce no or very little ellagic acid, respectively (Fig. 4B).

To confirm that the peroxidase activity present in the nectar is indeed due to MmNec3, we performed in-gel assays followed by proteomic analysis of activity bands. Proteins in raw nectar separated by native PAGE have peroxidase activity when using either guaiacol (general chromogenic substrate, Fig. 4C, left) or gallic acid (Fig. 4C, right) as a substrate. A small amount of iron was added to the substrate solution containing gallic acid to visualize ellagic acid formation in the gel; therefore, the dark activity band (Fig. 4C right) represents a complex of ellagic acid and Fe (HRP was used as a positive control). The activity band from the guaiacol-based assay was excised from the gel and subjected to proteomic analysis, which identified peptides covering 49.5% (97 of 196 amino acids) of the mature protein (Fig. 4D), thereby confirming that MmNec3 has peroxidase activity.

### In-vitro recapitulation of black nectar synthesis

We were next curious if a quasi-synthetic nectar could be made that recapitulates the color of the natural nectar. Starting with purified total nectar protein (Fig. 5A), successive additions of glucose, gallic acid, H_2_O_2_, and finally FeCl_3_ give a dark black product (Fig. 5A, fifth panel from left) with an absorbance spectrum nearly identical to that of raw nectar (Fig. 5B). A similar result is obtained if HRP is used as a positive control (Fig. 5B), but negative control reactions containing inactivated nectar protein do not (far right panel Fig. 5A and Fig. 5B). Cumulatively, these results strongly suggest that gallic acid is oxidized post-secretion by MmNec3 *in the nectar* to generate ellagic acid, which in turn complexes with near millimolar levels of Fe^3+^ to form a dark black color (Fig. 5C).

### Expression of genes potentially involved in black pigment formation

The ontological differences in nectar color suggested active control of nectar composition is occurring. Therefore, we examined global developmental and spatial expression in nectaries and leaves. *M. minor* flowers have well developed nectaries that are clearly distinct from surrounding tissues (Fig. S1C), making them easy to collect by manual dissection. Total RNA was isolated from floral nectaries at three developmental time points: (1) before secretion started (pre-secretory), (2) during active nectar secretion in open flowers at anthesis (secretory), and (3) in older open flowers with reduced nectar volume (post-secretory; all stages are further described in the *Materials & Methods*); RNA was also isolated from leaves as a tissue and spatial control. Bar-coded libraries from these four types of RNA (pre-secretory, secretory, post-secretory, and leaf) were made and sequenced via Illumina-based technology. These reads were assembled into contigs and mapped to the Arabidopsis genome by locus (Supporting File 1). The rationale for this approach is that Arabidopsis not only has the best annotated plant genome but is also the best studied in terms of nectary functional genomics (Bowman & Smyth, 1999; Baum *et al*., 2001; Lee *et al*., 2005a; Lee *et al*., 2005b; Kram & Carter, 2009; Bender *et al*., 2012; Bender *et al*., 2013; Lin *et al*., 2014; Wiesen *et al*., 2016). Collapsing all contigs, and associated reads, to individual Arabidopsis genes, allows for the most straight-forward identification of gene function as it relates to nectar/y biology.

*Melianthus* secretory nectaries express the typical suite of genes observed in those of other eudicots (Supp. File 1, Fig. 6). For instance, *SWEET9*, encoding a sucrose uniporter required for nectar secretion (Lin *et al*., 2014), has low expression in presecretory nectaries and leaves, but is strongly upregulated in secretory and post-secretory nectaries (Fig. 6). Conversely, leaves highly express genes required for photosynthesis (e.g., *RuBisCo* subunits, Fig. 6) but the nectaries do not. This result is not surprising since most nectaries do not photosynthesize. These results are indicative of high-quality sequence data that can be further mined for biologically relevant expression patterns.

The nectar peroxidase, *MmNec3*, is highly expressed in secretory and post-secretory nectaries, but its expression is very low in pre-secretory nectaries and leaves (Fig. 6). This developmentally- and spatially-enriched expression pattern suggests an important role for MmNec3 in secretory nectaries. Since the dark nectar at anthesis contains >10-fold more Fe than that of non-anthesis nectar (Fig. 3A), we also looked for the expression of genes involved in iron transport. In particular, apparent orthologs to Arabidopsis *ferritin 1* (*Fer1*) and *ferritin 2* (*Fer2*) are strongly induced in secretory over pre-secretory nectaries (Fig. 6). Plant ferritins play important roles in binding excess iron to limit oxidative stress and may also have a role in iron transport and storage (Kobayashi *et al*., 2019). Another candidate gene that might play a role in regulating Fe accumulation in nectar is annotated as a ‘heavy metal transport protein,’ which again is highly induced in secretory and post-secretory nectaries over pre-secretory nectaries and leaves (Fig. 6).

Nectar solutes, including proteins, are often synthesized de novo in nectaries as opposed to being secreted directly from the phloem (Carter, C & Thornburg, RW, 2004; Lin *et al*., 2014; Roy *et al*., 2017; Solhaug *et al*., 2019a; Solhaug *et al*., 2019b; Roy *et al*., 2022). Given the high levels of polyphenols in the nectar, we examined our transcriptomic data for the canonical gallic acid and galloyl glucose biosynthetic pathway in higher plants [as outlined in (Ye *et al*., 2022)]. Putative genes for each of the steps in galloyl glucose synthesis are highly expressed in pre-secretory nectaries relative to both secretory nectaries and leaves (Fig. S8). These results suggest that nectaries synthesize glucosylated polyphenols (i.e., gallotannins) de novo prior to export into the apoplast during nectar secretion. Interestingly, each of these steps is also upregulated in post-secretory nectaries over secretory nectaries, but the functional significance of this observation is unclear.

### Black nectar is visible and conspicuous to birds

Some colored nectars appear to serve as a visual cue to prospective pollinators, with the black nectars of *Melianthus* spp. possibly signaling the presence of a reward to passerine birds and perhaps bees (Hansen *et al*., 2007). To determine if this scenario is likely, we modeled the reflectance of the nectar, petals, and leaves from Fig. 1D onto the visual space of birds as well as bees. Since *M. minor* flowers are usually embedded inside the bush, leaf reflectance is assumed to be the primary background upon which floral reflectance (nectar and petals) resides. The vertices of the tetrahedron and the triangle shown in Fig. 7A,C correspond to different photoreceptors found in birds and bees, with reflectance data projected onto this visual space. We found that reflectance of *M. minor* flowers and nectar fall within the visual spaces of both birds and bees, confirming that birds as well as bees can indeed perceive the black nectar and red petals of *M. minor*. As per the avian visual model, chromatic contrast of the *M. minor* nectar is significantly higher than the achromatic contrast of the nectar and the chromatic and achromatic contrasts of the flowers (Fig. 7B; pairwise differences; all t > 3.00, p < 0.02; N = 36). As per the bee visual model however, achromatic contrast between flowers and leaves is higher than chromatic contrasts between flowers and leaves and the chromatic and achromatic contrasts between nectar and petals. However, this pattern is not statistically significant (Fig. 7D; all t < 2.5; all p > 0.05; N = 36). These differences in contrast values calculated using bird and bee visual models highlight how visual systems can shape how a specific object might appear vastly different to different animals. In the case of M. minor, while chromatic contrast between the nectar was the most conspicuous signal, achromatic contrast of petal is likely to be nectar of *M. minor* is a highly conspicuous signal produced by *M. minor* from a bird’s perspective but not bee’s perspective, therefore, likely to play an especially crucial role in attracting avian pollinators (Fig. 7).

## DISCUSSION

Our work addresses two overarching hypotheses: (1) that colored nectars contain novel pigment chemistries and (2) these pigments serve as visual attractants for prospective pollinators. With respect to the first hypothesis, it appears that almost all plant pigments are retained inside of cells [especially in vacuoles and plastids (Davies *et al*., 2022; Rodriguez-Concepcion & Daros, 2022)] instead of being exported. Therefore, extracellular pigments, like those found in colored nectars, are rare. Only ~70 plant species are known to produce colored nectars (Hansen *et al*., 2007), but they represent an opportunity to discover new colored compounds, as well as their synthesis and secretory pathways. Our study addresses the identity, synthesis, and biological function of the pigment that gives *Melianthus* nectar its jet-black color.

*Melianthus* nectar contains millimolar levels of polyphenols – like gallic acid, ellagic acid, and related compounds (Fig. 2) – which led us to hypothesize that it could be a natural analog of iron-gall ink. Relatively modern recipes for iron-gall ink [i.e., by governmental specification (Wailes, 1935)] call for the addition of gallic acid to FeSO4 (Fe^2+^; in combination with tannic acid), but we found that this mixture alone does not yield a black product (Fig. 3C). Conversely, mixtures of ellagic acid with FeCl_3_ (Fe^3+^) at biologically relevant concentrations immediately formed a black product that recapitulates the appearance and absorbance spectrum of the black nectar (Fig. 3C). The apparent discrepancy in the composition of iron-gall ink (rich in gallic and tannic acid) and that of black nectar (rich in ellagic acid and tannins) could be due to pH effects, as iron-gall ink is acidic [pH ~1-3 (Melo *et al*., 2022)] and the black anthesis nectar is alkaline (Fig. 1D). However, it is also well known that iron-gall inks darken significantly over time after exposure to air, which is thought to occur through the oxidation of Fe^2+^ to Fe^3+^ and associated complex formation with tannic acid (Lee *et al*., 2018). Based on our results, the darkening of iron-gall inks on paper also likely reflects the oxidation of gallic acid into ellagic acid *in situ*.

Ellagic acid is formed through the oxidative dimerization of gallic acid, which can be catalyzed by peroxidases in the presence of H_2_O_2_ (Kamel *et al*., 1977); however, the genes encoding the proteins involved in this process, and their associated *in vivo* biological functions, have not been reported. In this study, we identify a Type-III peroxidase, MmNec3, secreted into the nectar that can oxidize gallic acid to form ellagic acid (Fig. 4). Given the high levels of free gallic acid in the nectar (Fig. 2), the most parsimonious explanation for the role of MmNec3 in the nectar is to catalyze the conversion of gallic acid into ellagic acid to support pigment formation. This supposition is reinforced by the finding of significantly higher levels of ellagic acid and lower levels of gallic acid in anthesis nectar relative to non-anthesis nectar (Fig. 2C) and that relatively high levels of ellagic acid compared to gallic acid are found in the nectars of multiple *Melianthus* species (Fig. S3). Intriguingly, the involvement of nectar enzymes in pigment formation is not unique, as three proteins in the red nectar of *Nesocodon mauritianus* are either necessary or sufficient for the formation of a red colored alkaloid (Roy *et al*., 2022).

Several key questions remain on how the black pigment forms in the nectar. First, in addition to free gallic and ellagic acid, *Melianthus* nectar contains gallo- and ellagitannins (i.e., hydrolysable tannins; Fig. 2B, Fig. S3, Table S1). It is unclear if gallic acid is secreted into nectar as a free monomer or as a glucosylated conjugate (gallotannin), but the latter situation would necessitate the presence of a glucosyl hydrolase to release gallic acid from glucose to achieve the high levels of free gallic and ellagic acid observed in *Melianthus* nectar. Second, the source of H_2_O_2_ needed for peroxidase activity is also unknown, but nectars appear to commonly contain mechanisms for H_2_O_2_ production (Carter & Thornburg, 2000; Carter, C & Thornburg, RW, 2004; Carter, CJ & Thornburg, RW, 2004; Roy *et al*., 2022). Third, anthesis nectar contains very high levels of iron (~0.7 mM, Fig. 3A), which is required for pigment formation (Fig. 3B-D) through complexes with free ellagic acid. The mechanism through which Fe is exported into the nectar is unclear, but several genes, like ferritins and a putative heavy metal transport protein (Fig. 6), are promising candidates that need further investigation. Lastly, it is unknown how the anthesis nectar becomes alkaline, but pH does not appear to greatly impact color intensity (Fig. S5).

*Melianthus* flowers are almost exclusively visited by avian pollinators (Scott-Elliot & Cantab, 1890; Henning, 2003; Pauw & Stanway, 2015), in contrast to the most closely related genus (*Bersama*), which is visited by a range of insect pollinators (Verdcourt, 1956). Our results are consistent with the hypothesis that the black nectar of *M. minor* is conspicuous to birds, especially relative to bees (Fig. 7). Our analyses focused on the contrast of the black nectar with the red petals of the flowers, for a pollinator viewing the flower head-on, but it is important to note that the nectar is also visible through the translucent sepals of the flower (Fig. 1, Fig. S1, and Henning 2003). These observations, coupled with previous information on the natural and evolutionary history of this genus (Fig. 8, Henning 2003, Linder *et al*. 2006) suggest a few evolutionary scenarios that could explain the origin and function of the black nectar in *Melianthus*. First, previous studies have noted that patterns of nectar coloration across the phylogeny of *Melianthus* suggest three separate origins of black nectar (Fig. 8). The present results add to this broader context with the observation that variation in nectar coloration can emerge based on variation in the chemistry of the nectar components (Fig. 3). In other words, the variation observed across species of *Melianthus* could potentially be due to variations in the timing of nectar sampling and observations, and not independent evolutionary origins. In existing studies, it is unclear how frequently, and when in the life course of a flower, previous nectar observations were made (Henning 2003); clearly, we need more nectar observations and chemical studies across different species of *Melianthus* to reconstruct the evolutionary history of the black nectar. Second, our results are consistent with the potential role of black nectar being visually conspicuous to avian pollinators. However, existing phylogenetic work suggests that black nectar evolved after the shift to avian pollinators (Fig. 8), calling into question whether the attraction of pollinators per se was the primary selective factor in its origins. Additional natural history observations show that black nectar is associated with species with smaller volumes of nectar and species in more arid conditions (Fig. 8, Henning 2003). Interestingly, the morphology of flowers in the *M. minor* clade has adaptations associated with reducing the evaporative loss of nectar in dry climates (e.g., “hiding” the nectar), likely a significant issue for bird-pollinated plants which produce large volumes of nectar. It is possible that the black coloration of nectar, coupled with transparent sepals, could act as a signal to pollinators while shielding nectar from evaporation, a potential benefit to both the pollinator and plant. Future experimental studies across species of *Melianthus* could address this hypothesis by looking at nectar evaporation rates and avian attraction to “hidden” nectar of different colors (non-colored, gray, black).

**Fig. 8.**
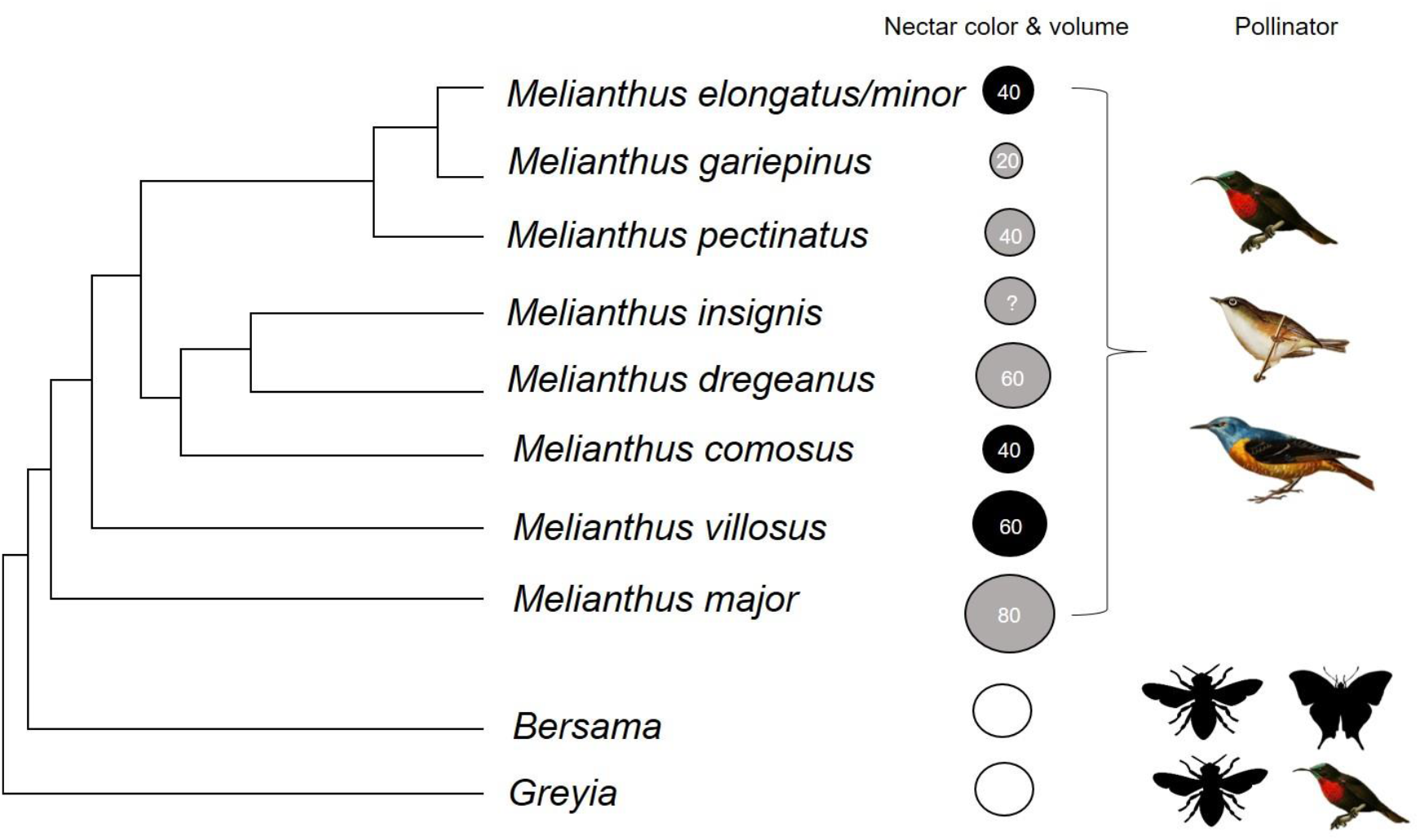
Phylogenetic tree of *Melianthus* spp. showing nectar properties and associated pollinators. Nectar volume (μL) and color according to Henning (2003) are indicated in the circles, aligned with a molecular phylogeny (Henning 2003, Linder *et al*. 2006). Reported pollinators are also indicated. All images have a creative commons license.

Our study focused on a role for the black nectar serving as a visual cue to pollinators, but we cannot discount other non-mutually exclusive functions not related to its color. For instance, the polyphenolics may have antimicrobial activity and just happen to be colored. Similarly, the nectar pigment may be distasteful to some animals and serve as a filter to prevent unwanted visitors who may otherwise consume the nectar without providing pollination services (Johnson *et al*., 2006). Anecdotally, we did not observe any unpleasant flavor in Melianthus nectar, but additional studies are needed to elucidate the black color’s true biological function.

## DATA AVAILABILITY

RNA-seq data for *Melianthus* nectaries and leaf tissues are available at NCBI SRA via accession # (pending). The sequence of MmNec3 is available on NCBI GenBank via accession #OQ077505. The mass spectrometry proteomics data have been deposited to the ProteomeXchange Consortium via the PRIDE (Perez-Riverol *et al*., 2022) partner repository with the dataset identifiers PXD038938 and PXD038940 with DOIs of 10.6019/PXD038938 and 10.6019/PXD038940. Raw data for all other figures and tables is available upon request.

## ACKNOWLEDGEMENTS

The authors recognize the Center for Mass Spectrometry and Proteomics at the University of Minnesota and various supporting agencies, including the NSF for Major Research Instrumentation grants 9871237 and NSF-DBI-0215759 used to purchase the instruments described in this study. We thank the University of Minnesota Genomics Center for conducting RNA quality control, RNA-seq library creation, next-generation sequencing, and primary NGS data QC. We also thank Dr. Rachel Vannette, Dino Sbardellati, Shawn Christensen, Jacob Francis, and Ernesto Sandoval, all from the University of California Davis, for providing access to *Melianthus* materials and additional research support, including extremely helpful sample collection and access to laboratory resources. Lastly, we thank Ruth Cozien of the Center for Functional Biodiversity, School of Life Sciences, University of Kwa-Zulu Natal in South Africa for generously providing nectar samples collected from *M. pectinatus* and *M. villosus*. This work was supported by grants from the US NSF to M.H. and C.J.C. (IOS-1339246); E.C.S.-R., M.H., A.D.H., and C.J.C. (IOS-2025297); and A.D.H. (IOS-1238812). R.R. was supported by a postdoctoral fellowship from the United States Department of Agriculture (2018-67012-28038).

## AUTHOR CONTRIBUTIONS

EM, RR, AZ, MH, ESR, AH and CC conceived the experimental approach. EM, RR, KFS, AZ, AH, KB and CC conducted the experiments. MH and AH conducted a majority of the informatics analyses of RNA-seq and metabolomics data, respectively, with additional support from EM and CC. AZ collected reflectance data and conducted visual response modeling. ESR placed the results within the context of *Melitanthus* spp. natural history and evolution. EM, ESR, and CC wrote a majority of the manuscript with contributions from all co-authors.

## SUPPORTING INFORMATION

Supporting Figures – Fig. S1 - S8 and Table S1

Supporting File 1 – summarized RNA-seq data of nectaries and leaves, including annotations, normalized read counts, and statistical analyses

Supporting File 2 – nucleotide sequences of contigs from RNA-seq data

Supporting File 3 – translated contig sequences

