## Supplementary material for "Post-secretory synthesis of a natural analog of iron-gall ink in the black nectar of *Melianthus* spp.": Supp file 1 - Figs S1-S8 and Table S1

**This PDF file includes:**

Figures S1 to S6 and Table S1

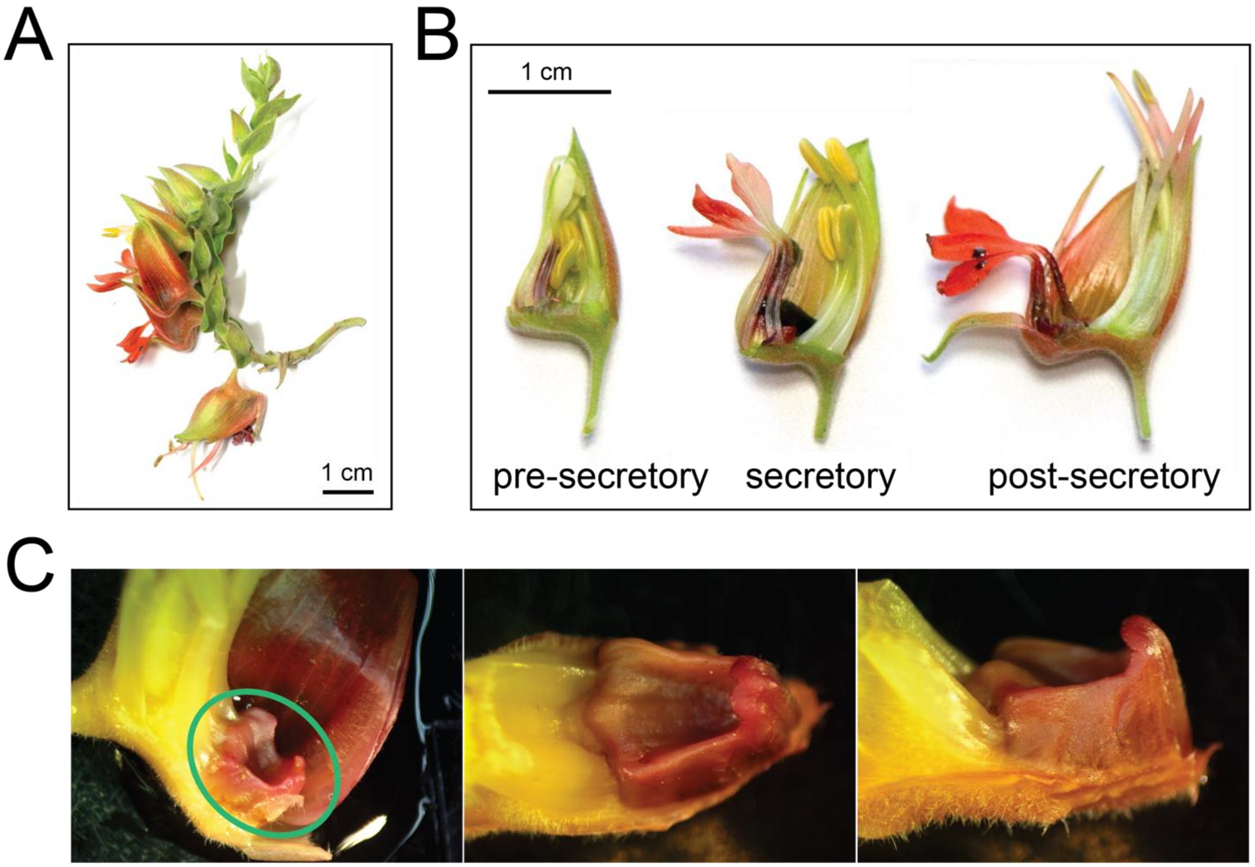


**Fig. S1. *M. minor* floral structures.** (A) Inflorescence of *M. minor* displaying the various phases of flower development. (B) Longitudinal cross-sections of flowers at the pre-secretory, secretory, and post-secretory time points. These phases of development correspond to those utilized in the transcriptome analysis. (C) The nectary in *M. minor*. The nectary is circled by a green oval in the first image. The subsequent images highlight the morphology of the nectary. It is distinguished by its bright, contrasting red coloration.

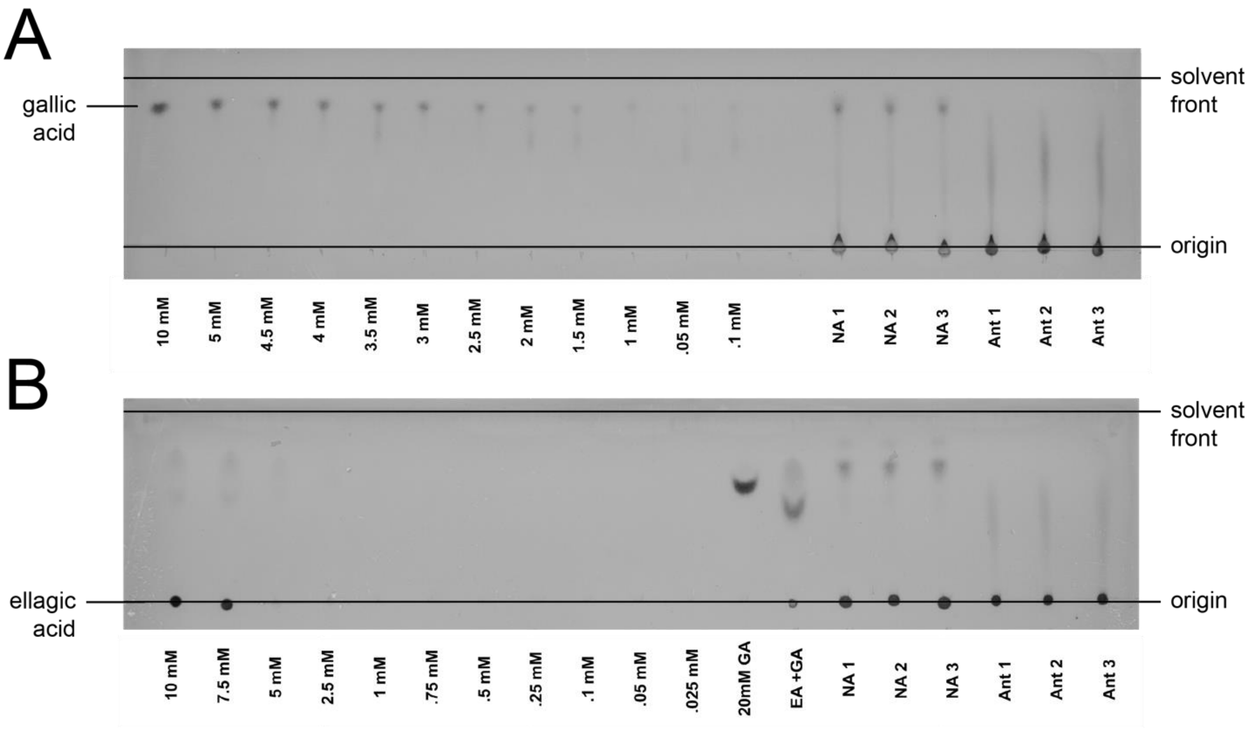


**Fig. S2. Gallic acid and ellagic acid quantification.** (A) The greyscale version of the TLC plate used to measure gallic acid in non-anthesis and anthesis samples. The concentrations are noted at the bottom of each samples. NA = non-anthesis. Ant = anthesis. Triplicates of each sample type were used (B) TLC plate used to calculate the concentration of ellagic acid in triplicates of non-anthesis (NA) and anthesis (Ant) nectar. The origin and solvent lines are denoted in the figure. Ellagic acid, under these circumstances, is immobile, while gallic acid is mobile.

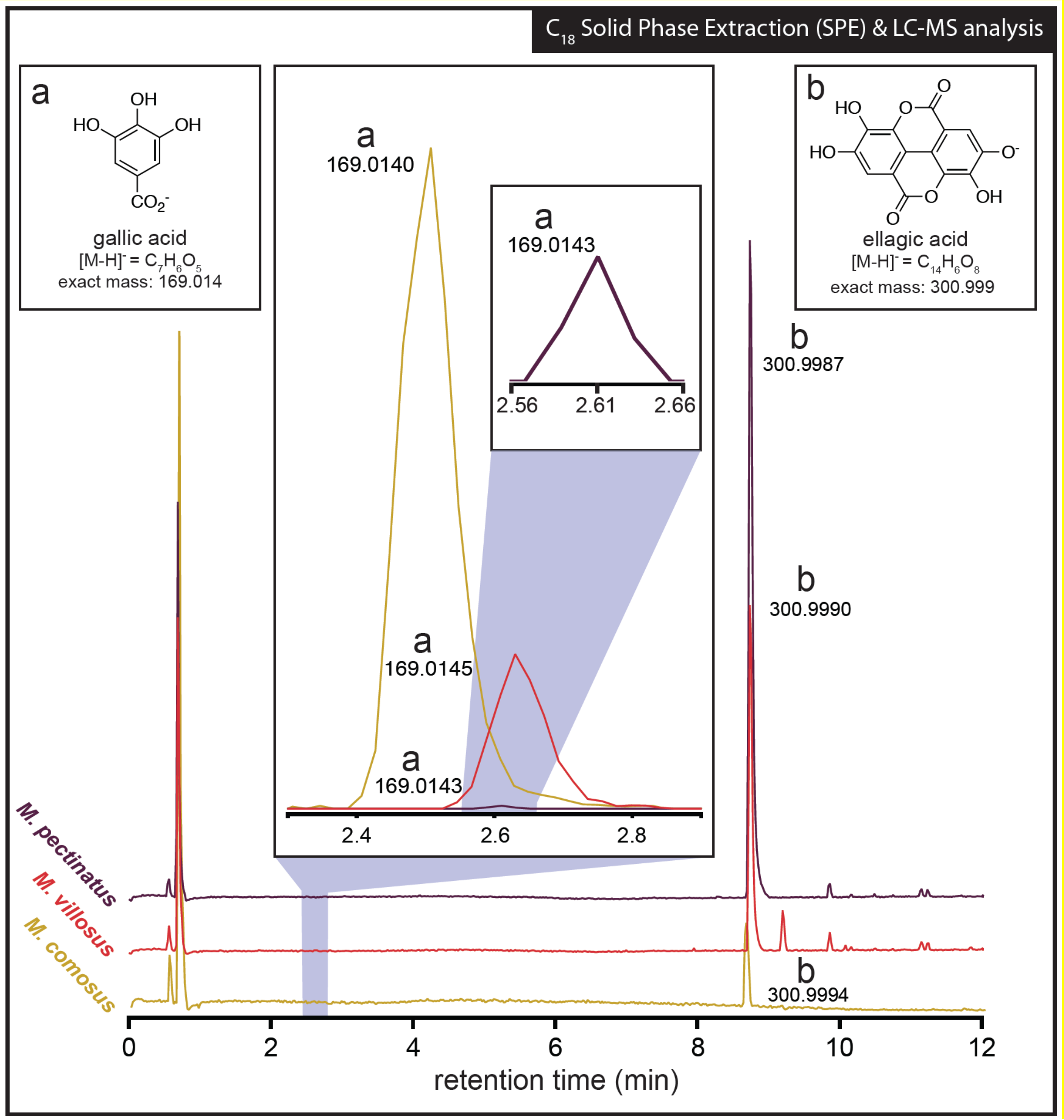


**Fig. S3. *Melianthus comosus, M. pectinatus, and M. villosus* nectars also contain gallic and ellagic acid.** The LC MS profiles for gallic and ellagic acid in these nectars are shown, which were analyzed in the same manner as for *M. minor* nectar shown in Fig. 2A.

**Table S1.** Tannins and polyphenols found in *Melianthus* spp. nectars

| | compound | *M. comosus* | *M. minor* | *M. pectinatus* | *M. villosus* |
|---|---|---|---|---|---|
| 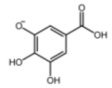 | gallic acid<br>[M-H] = $C_7H_6O_5$<br>169.014 | + | + | + | + |
| 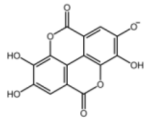 | ellagic acid<br>[M-H] = $C_{14}H_6O_8$<br>300.999 | + | + | + | + |
| 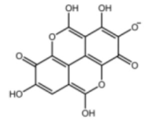 | flavellagic acid<br>[M-H] = $C_{14}H_5O_9^-$<br>316.9939 | + | + | + | + |
| 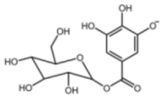 | monogalloyl glucose<br>[M-H] = $C_{13}H_{15}O_{10}^-$<br>331.0671 | + | + | + | + |
| 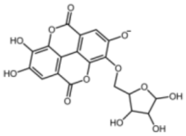 | monogalloyl pentose<br>[M-H] = $C_{19}H_{13}O_{12}^-$<br>433.0412 | + | + | + | + |
| 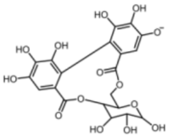 | monoellagoyl glucose<br>[M-H] = $C_{20}H_{17}O_{14}^-$<br>481.0624 | — | + | — | — |
| 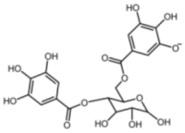 | digalloyl glucose<br>[M-H] = $C_{20}H_{19}O_{14}^-$<br>483.0780 | + | + | — | + |
| 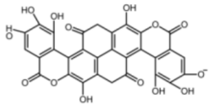 | gallagic acid dilactone<br>[M-H] = $C_{28}H_9O_{16}^-$<br>600.9896 | — | — | + | + |
| 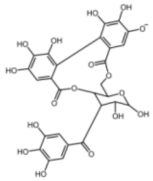 | monogalloyl-<br>monoellagoyl glucose<br>[M-H] = $C_{27}H_{21}O_{18}^-$<br>633.0733 | — | + | — | — |
| 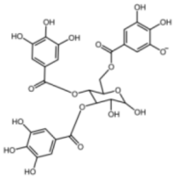 | trigalloyl glucose<br>[M-H] = $C_{27}H_{23}O_{18}^-$<br>635.0890 | + | + | — | + |
| 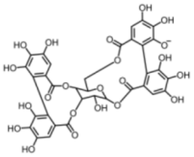 | diellagoyl glucose<br>[M-H] = $C_{34}H_{23}O_{22}^-$<br>783.0686 | + | — | — | — |
| 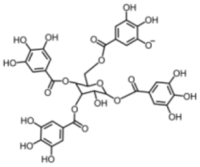 | tetragalloyl glucose<br>[M-H] = $C_{34}H_{27}O_{22}^-$<br>787.0999 | + | — | — | + |

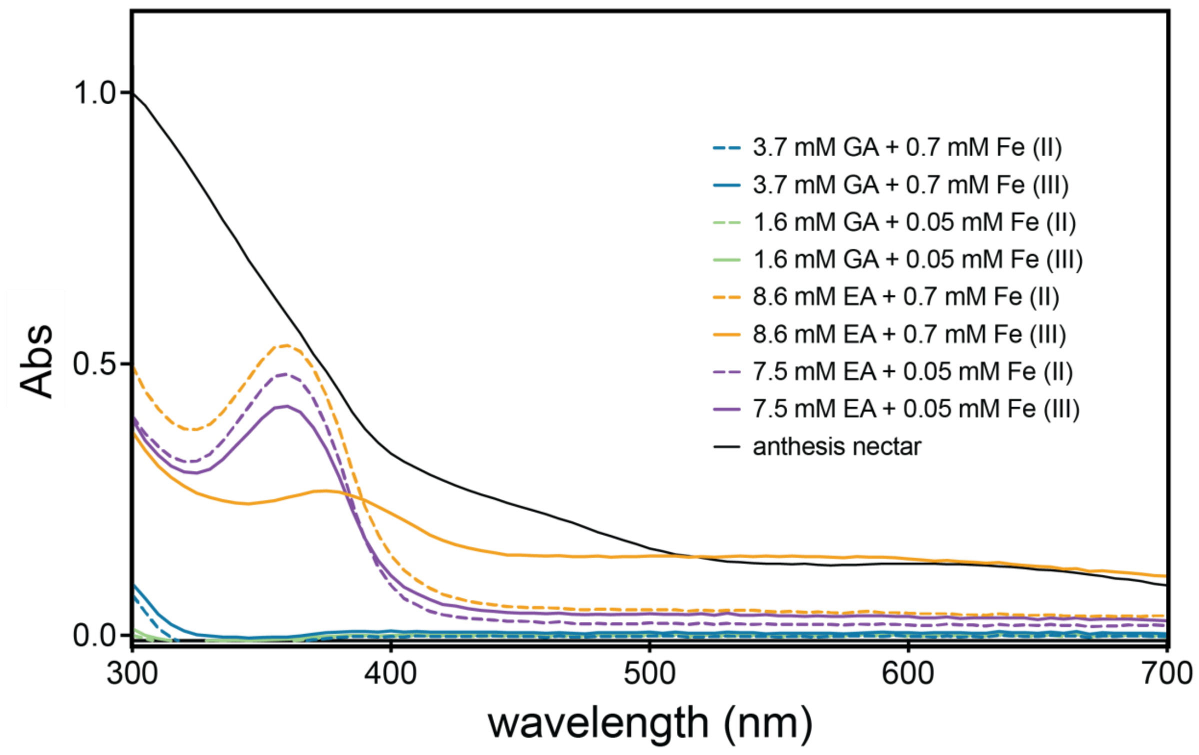


**Fig. S4. Ferrous and ferric iron addition to gallic acid and ellagic acid.** (A) The absorbance spectra of varying concentrations of both forms of iron ($Fe^{2+}$ and $Fe^{3+}$) to biologically relevant levels of gallic acid and ellagic acid in pH 8.5 buffer (25 mM HEPES). Anthesis nectar was diluted 1:20 using sterile water, while the rest of the samples were diluted 1:10 with sterile water.

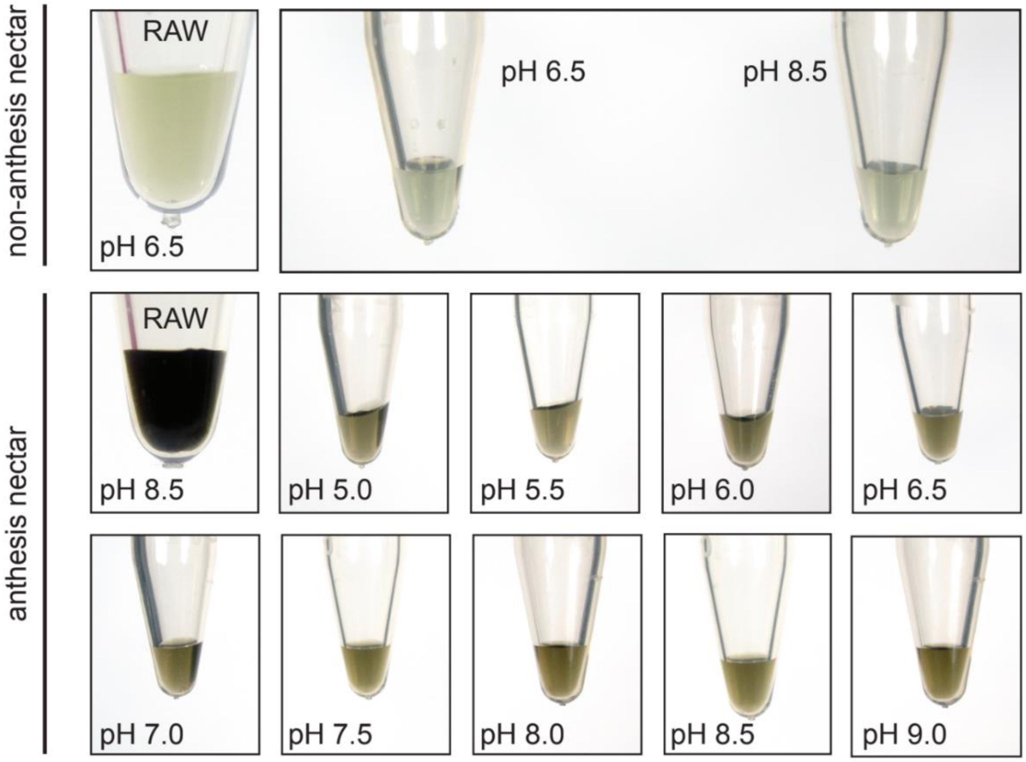


**Fig. S5. Influence of pH on nectar color intensity.** Non-anthesis and anthesis nectar samples were diluted 1:3 in buffers of varying pH, with the resulting appearance of the nectar being shown. The buffers used were all 50 mM, including sodium acetate (pH 5.0 and 5.5), MES (pH 6.0 and 6.5), HEPES (pH 7.0, 7.5, 8.0) and Tricine (pH 8.5 and 9.0).

### MmNec3

| | | | | | |
|---|---|---|---|---|---|
| MSSLHQLLLL | SFLLLPFSYA | DLRLGFYR**SS** | **CPQAESIVQQ** | **VVR**ERFNR**DR** | 50 |
| **SITAAFLR**MH | FHDCFVSGCD | ASILIDATAG | KPSEK**NAGPN** | **LTVRGNEVID** | 100 |
| **EIKRRLEQAC** | **PSTVSCADII** | **TLATRDSVVL** | **SGGPNYDVPT** | **GR**QDGLVSNP | 150 |
| DNVNLPGPTL | SVSQALQSFT | EKGFNLAEMV | TLLGAHSVGF | SHCSFFQDR**L** | 200 |
| **TDFQGTGAPD** | **RNMDPALVTR** | **LRDTCAQPND** | **PTVFLDQGTP** | **NAMDNKWYSQ** | 250 |
| **LIQKRGILQI** | **DQALAIDRSS** | **ASIVSGFASN** | **DASFR**QR**FAA** | **AMIKMGSLQL** | 300 |
| **NGPGEIRK**NC | RAFNKA | | | | 316 |

**Fig. S6. Amino acid sequence of MmNec3 and peptides identified from proteomic analyses of the band shown in Fig. 4A.** The amino acids underlined at the N-terminus reflect the predicted N-terminal signal peptide for secretion from the cell. Bolded regions correspond to tryptic peptides found LC MS/MS.

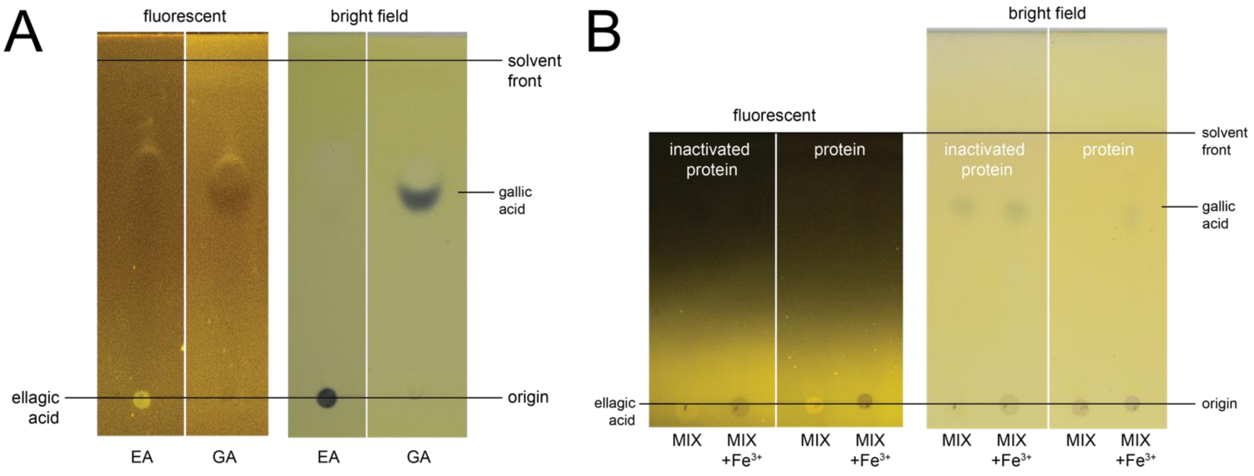


**Fig. S7. Conversion of gallic acid to ellagic acid by nectar peroxidase activity.** (A) Gallic acid and ellagic acid standards demonstrating that ellagic acid is fluorescent under these conditions, and that gallic acid reacts with the iron ($Fe^{3+}$) to produce a noticeable black band. Ellagic acid (EA) is in the left panel of both the fluorescent and bright field images. Gallic Acid (GA) is in the right panel of both images. (B) Inactivated proteins (Proteinase K Solution) and purified proteins were used to judge the ability of MmNec3 to convert gallic acid to ellagic acid in the presence of $H_2O_2$. MIX (in HEPES pH 8.0) = protein + gallic acid + glucose + $H_2O_2$. The samples to the right of the MIX have $Fe^{3+}$ [0.7 mM] added to them prior to TLC. Again, ellagic acid is fluorescent at the baseline, while gallic acid is mobile and forms a black colorant when complexed with the iron spray. Note the decrease of gallic acid in reactions containing active protein in the far right panel.

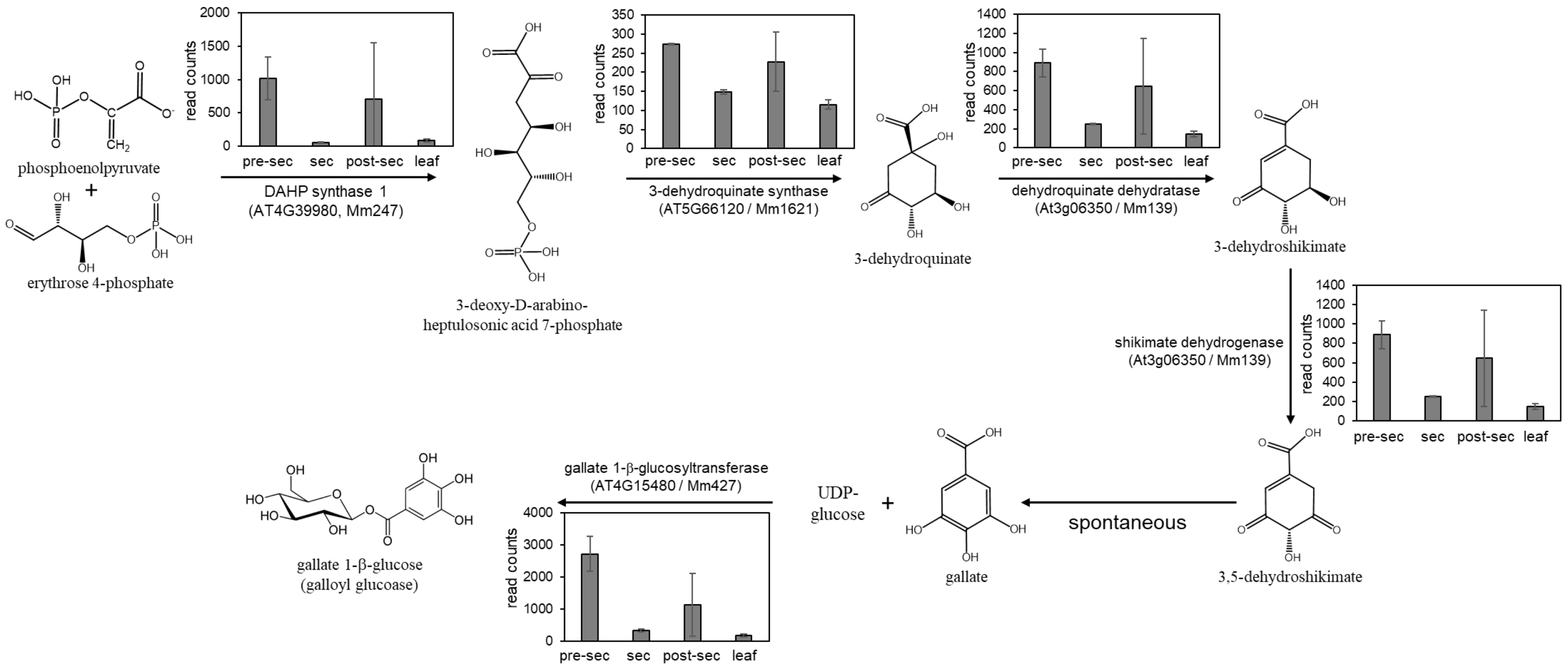


**Fig. S8. Expression of genes putatively involved in gallic acid and galloyl glucose biosynthesis**. Normalized read counts derived from RNA-seq analysis are shown for select genes in pre-secretory, secretory, and post-secretory nectaries, as well as leaves. Each panel indicates the putative identity along with the contig number (e.g., Mm247) and top Arabidopsis hit (e.g., At4g39980). Triplicate samples were used for all tissues except for pre-secretory nectaries, which only had two samples. Error bars represent standard deviation. Summarized data are shown in Supplemental File 1 and raw data is available on NCBI SRA via accession # (pending). This pathway is based on that described in Ye et al. *Molecules* **2022**, *27*(23), 8374; **https://doi.org/10.3390/molecules27238374**
